# Trajectory-based differential expression analysis for single-cell sequencing data

**DOI:** 10.1101/623397

**Authors:** Koen Van den Berge, Hector Roux de Bézieux, Kelly Street, Wouter Saelens, Robrecht Cannoodt, Yvan Saeys, Sandrine Dudoit, Lieven Clement

**Affiliations:** Department of Applied Mathematics, Computer Science and Statistics, Ghent University, Ghent, Belgium; Bioinformatics Institute Ghent, Ghent University, Ghent, Belgium; Division of Biostatistics, School of Public Health, University of California, Berkeley, CA, USA; Department of Data Sciences, Dana-Farber Cancer Institute, Boston, MA, USA; Data mining and Modelling for Biomedicine, VIB Center for Inflammation Research, Ghent, Belgium; Center for Medical Genetics, Ghent University Hospital, Ghent, Belgium; Department of Biomolecular Medicine, Ghent University, Ghent, Belgium; Department of Statistics, University of California, Berkeley, CA, USA

## Abstract

Trajectory inference has radically enhanced single-cell RNA-seq research by enabling the study of dynamic changes in gene expression levels during biological processes such as the cell cycle, cell type differentiation, and cellular activation. Downstream of trajectory inference, it is vital to discover genes that are associated with the lineages in the trajectory to illuminate the underlying biological processes. Furthermore, genes that are differentially expressed between developmental/activational lineages might be highly relevant to further unravel the system under study. Current data analysis procedures, however, typically cluster cells and assess differential expression between the clusters, which fails to exploit the continuous resolution provided by trajectory inference to its full potential. The few available non-cluster-based methods only assess broad differences in gene expression between lineages, hence failing to pinpoint the exact types of divergence. We introduce a powerful generalized additive model framework based on the negative binomial distribution that allows flexible inference of (i) within-lineage differential expression by detecting associations between gene expression and pseudotime over an entire lineage or by comparing gene expression between points/regions within the lineage and (ii) between-lineage differential expression by comparing gene expression between lineages over the entire lineages or at specific points/regions. By incorporating observation-level weights, the model additionally allows to account for zero inflation, commonly observed in single-cell RNA-seq data from full-length protocols. We evaluate the method on simulated and real datasets from droplet-based and full-length protocols, and show that the flexible inference framework is capable of yielding biological insights through a clear interpretation of the data.

---

Single-cell RNA sequencing (scRNA-seq) has revolutionized modern biology by allowing researchers to profile transcript abundance at the resolution of single cells. This has opened new avenues to study cellular pathways during cell cycle, cell type differentiation, or cellular activation. Indeed, scRNA-seq can provide a snapshot of the transcriptomes of thousands of single cells in a cell population, which are each at distinct points of the dynamic process under study. This wealth of transcriptional information, however, presents many data analysis challenges. Until recently, statistical and computational efforts have focused mostly on trajectory inference (TI) methods, which aim to first allocate cells to lineages and then order them based on pseudotimes within these lineages. A wide range of TI methods have been proposed; 45 of which are extensively benchmarked in Saelens et al. [2019]. Note that we use the term *trajectory* to refer to the collection of *lineages* for the process under study.

Most TI methods share a common workflow: dimensionality reduction followed by inference of lineages and pseudotimes in the reduced-dimensional space [Cannoodt et al., 2016]. While early methods were limited to inferring trajectories comprised of a single linear lineage, recent developments have allowed the inference of trajectories that might bifurcate multiple times and consist of several smooth lineages, or that might have cyclic patterns [Street et al., 2018, Lönnberg et al., 2017, Qiu et al., 2017]. These advances in TI methods enable researchers to study dynamic biological processes, such as complex differentiation patterns from a progenitor population to multiple differentiated cellular states [Byrnes et al., 2018, Herring et al., 2018], and have the promise to provide transcriptome-wide insights into these processes.

Unfortunately, statistical inference methods are lacking to identify genes associated with lineage differentiation and to unravel how their corresponding transcriptional profiles are driving the dynamic processes under study. Indeed, differential expression (DE) analysis of individual genes along lineages is often performed on discrete groups of cells in the developmental pathway, e.g., by comparing clusters of cells along the trajectory or clusters of differentiated cell types. Such discrete DE approaches do not exploit the continuous expression resolution that can be obtained from the pseudotemporal ordering of cells along lineages provided by TI methods. Moreover, comparing cell clusters within or between lineages can obscure interpretation: it is often unclear which clusters should be compared, how to properly combine the results of several pairwise cluster comparisons, or how to account for the fact that not all these comparisons are independent of each other. Inevitably, the number of cluster comparisons also increases rapidly with the number of lineages of interest, leading to multiple testing issues at the gene level [Van den Berge et al., 2017] and further decreasing the reproducibility of scRNA-seq DE results.

A few methods have been been published with the aim of improving trajectory-based differential expression analysis by modeling gene expression as a smooth function of pseudotime along lineages. `GPfates` [Lönnberg et al., 2017] relies on a mixture of overlapping Gaussian processes [Lázaro-Gredilla et al., 2012], where each component of the mixture model represents a different lineage. For each gene, the method tests whether a model with a bifurcation significantly increases the likelihood of the data as compared to a model without a bifurcation, essentially testing whether gene expression is differentially associated with the two lineages. Similarly, the `BEAM` approach in `Monocle 2` [Qiu et al., 2017] allows users to test whether differences in gene expression are associated with particular branching events on the trajectory. Both methods improve upon a discrete cluster-based approaches by: (1) exploiting the continuous expression resolution along the trajectory and (2) comparing lineages using a single test based on entire gene expression profiles. However, they both lack interpretability, as they cannot pinpoint the regions of the gene expression profiles that are responsible for the differences in expression between lineages. Moreover, the `GPfates` model is restricted to trajectories consisting of just one bifurcation, essentially precluding its application to biological systems with more than two lineages (i.e., a multifurcation or more than one bifurcation). `BEAM` is restricted to the few dimensionality reduction methods that are implemented in the `Monocle` software, namely, independent component analysis (ICA), DDRTree [Qiu et al., 2017], and uniform manifold approximation and projection (UMAP) [McInnes et al., 2018]. Hence, novel methods to infer differences in gene expression patterns within or between transcriptional lineages with complex branching patterns are vital to further advance the field.

In this manuscript, we introduce `tradeSeq`, a method and software package for trajectory-based differential expression analysis for sequencing data. `tradeSeq` provides a flexible framework that can be used downstream of any dimensionality reduction and TI methods. Unlike previously proposed approaches, `tradeSeq` provides several tests that each identify a distinct type of differential expression pattern along a lineage or between lineages, leading to clear interpretation of the results. In practice, `tradeSeq` infers smooth functions for the gene expression measures along pseudotime for each lineage using generalized additive models and tests biologically meaningful hypotheses based on parameters of these smoothers. By allowing cell-level weights for each individual count in the gene-by-cell expression matrix, `tradeSeq` can handle zero inflation, which is essential for dealing with dropouts in full-length scRNA-seq protocols [Van den Berge et al., 2018]. As it is agnostic to the dimensionality reduction and TI methods, the approach scales from simple to complex trajectories with multiple bifurcations: `tradeSeq` only requires the original expression count matrix of the individual cells, estimated pseudotimes, and a hard or soft assignment (weights) of the cells to the lineages to infer the lineage-specific smoothers. For within-lineage differential expression, `tradeSeq` provides both global tests to screen for genes with overall DE along a lineage, as well as specific tests to pinpoint relevant variation in gene expression profiles within the lineage. Likewise, for between-lineage comparisons, `tradeSeq` provides both global tests to compare expression patterns between entire lineages (useful for initial screening of interesting genes), as well as specific tests that allow researchers to pinpoint relevant differences in expression profiles between lineages. If multiple hypotheses are assessed for each gene, one can build upon our `stageR` package [Van den Berge et al., 2017] to conduct an omnibus test (e.g., there are no differences in expression profiles across multiple lineages) prior to *post hoc* tests that identify the relevant specific differences (e.g., all pairwise comparisons between lineages). We benchmark our method against current state-of-the-art methods using simulated datasets (with cyclic, bifurcating, and multifurcating trajectories) and demonstrate its functionality and versatility on two real datasets. These case studies highlight the enhanced interpretability of `tradeSeq`’s results, which lead to improved understanding of the underlying biology.

## Methods

In this Section, we first present a negative binomial generalized additive model for expression measures along a trajectory. Building on this model, we then describe a general and flexible framework for identifying genes that are differentially expressed either within or between lineages of a given trajectory.

### Negative binomial generalized additive models

We build on the generalized additive model (GAM) methodology to model gene expression profiles as non-linear functions of pseudotime for the different lineages in a complex trajectory. In our GAM framework, each lineage is represented by a separate cubic smoothing spline, i.e., a linear combination of cubic basis functions of pseudotime. The flexibility of GAM also allows us to easily adjust for other covariates or confounders such as treatment and batch. The discrete nature and the over-dispersion of read counts is addressed by modeling the expression measures *Y*_*gi*_, for a given gene *g* ∈{1,…, *G*} across cells *i ∈* {1,…, *n*}, using a negative binomial (NB) distribution with cell and gene-specific means *µ*_*gi*_ and gene-specific dispersion parameters *ø*_*g*_. Hence, we propose the following gene-wise negative binomial generalized additive model (NB-GAM)

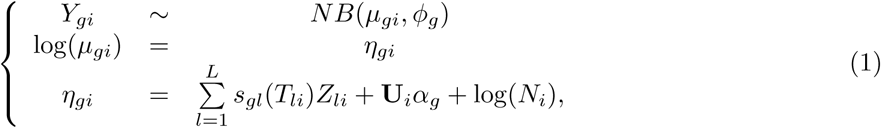

where the mean *µ*_*gi*_ of the NB distribution is linked to the additive predictor *η*_*gi*_ using a logarithmic link function. The gene-wise additive predictor consists of lineage-specific smoothing splines *s*_*gl*_, that are functions of pseudotime *T*_*li*_, for lineages *l* ∈{1,…, *L*}. The binary matrix Z = (*Z*_*li*_ ∈{0, 1}: *l* ∈{1,…, *L*}, *i* ∈ {1,…, *n*}) assigns every cell to a particular lineage based on user-supplied weights (e.g., from `slingshot` [Street et al., 2018] or `GPfates` [Lönnberg et al., 2017], see details in Supplementary Methods). We let *ℒ*_*l*_ ={*i*: *Z*_*li*_ = 1} denote the set of cells assigned to lineage *l*. In addition, we allow the inclusion of *p* known cell-level covariates (e.g., batch, age, or gender), represented by an *n* × *p* matrix U, with *i*^th^ row U_*i*_ corresponding to the *i*^th^ cell, and a regression parameter *α*_*g*_ of dimension *p×* 1. Differences in sequencing depth or capture efficiency between cells are accounted for by cell-specific offsets *N*_*i*_.

The smoothing spline *s*_*gl*_, for a given gene *g* and lineage *l*, can be represented as a linear combination of *K* cubic basis functions,

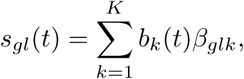

where the cubic basis functions *b*_*k*_(*t*) are enforced to be the same for all genes and lineages. In our implementation, we set *K* = 10. Thus, for each gene and each lineage in the trajectory, we estimate *K* = 10 regression coefficients *β*_*glk*_. The number of parameters *L ×K* + *p* + 1 in the gene-wise model is therefore typically much lower than the number of cells *n* in the dataset. In practice, we found the results to be robust to the choice of *K*. Indeed, the proportion of deviance explained by the model is not altered for any given choice of *K* between 6 and 14 (Supplementary Figure S1).

The NB-GAM is fitted gene by gene using the `fitGAM` function from the `tradeSeq` package, which relies on the `mgcv` package in R. We build upon recent developments that allow the joint estimation of the NB regression parameters in *µ*_*gi*_ and of the dispersion parameter *Ø*_*g*_ [Wood et al., 2016]. In order to control the smoothness of the spline, some coefficients *β*_*glk*_ are shrunken by substracting a penalty 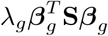 from the log-likelihood function, where *β*_*g*_ denotes the concatenation of the *L K*-dimensional column vectors *β*_*gl*_ of lineage-specific smoother coefficients and S is an (*LK*) *×* (*LK*) diagonal matrix that indicates which coefficients in *β*_*g*_ are to be penalized. The magnitude of penalization is controlled by the smoothing parameter *λ*_*g*_, which is selected using generalized cross-validation [Wood, 2017]. Note that we enforce identical basis functions between lineages, i.e., *b*_*k*_ does not depend on *l*, as well as identical smoothing parameter *λ*_*g*_, in order to ensure that the smoothers are comparable across lineages.

Importantly, the model of Equation (1) can accommodate zero-inflated counts typical for full-length scRNA-seq protocols by using observation-level (i.e., cell-level) weights obtained from the zero-inflated negative binomial (ZINB) approach of Van den Berge et al. [2018] and Risso et al. [2018a].

## Statistical inference

We propose a general and flexible testing framework for (linear combinations of) the parameters *β*_*g*_, which allows us to pinpoint specific types of differences in gene expression both within and between lineages, see Figure 1 for an overview. We first present the general approach and then detail the implementation and interpretation of specific DE tests.

**Figure 1:**
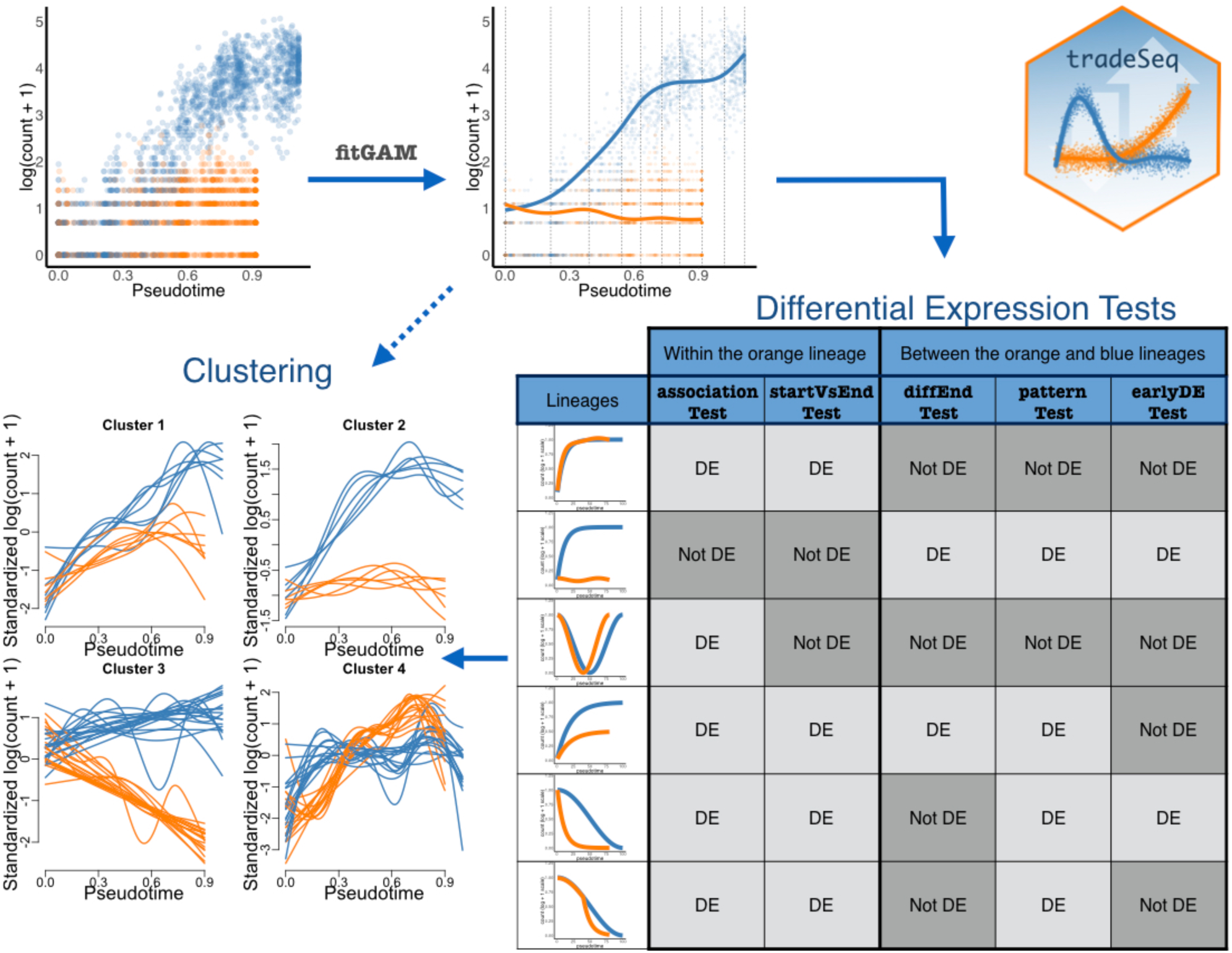
Overview of tradeSeq functionality. For each gene, we start with a scatterplot of expression measures vs. pseudotimes, where each lineage is represented by a different color (top left). A NB-GAM is fitted using the `fitGAM` function. The locations of the knots for the splines are displayed with gray dashed lines. The NB-GAM can then be used to perform a variety of tests of differential expression within or between lineages, as well as for clustering of the expression profiles of DE genes. In the table, we assume that the `earlyDETest` is used to assess differences in expression patterns early in the lineage, e.g., with option `knots = c(1, 2)`.

All proposed DE procedures involve testing null hypotheses of the form *H*_0_: C^*T*^ *β*_*g*_ = 0 using Wald test statistics

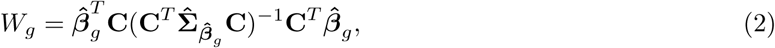

where 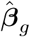 denotes an estimator of *β*_*g*_, 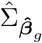 represents an estimator of the covariance matrix 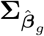 of 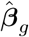, and C is an (*LK*) ×*C* matrix representing the *C* contrasts of interest for the DE test.

For each gene, we compute *p*-values based on the nominal chi-squared asymptotic null distribution of the Wald statistics (with degrees of freedom equal to the column-rank of C). Rather than attaching strong probabilistic interpretations to the *p*-values (which, as in most RNA-seq applications, would involve a variety of hard-to-verify assumptions and would not necessarily add much value to the analysis), we view the *p*-values simply as useful numerical summaries for ranking the genes for further inspection.

### Within-lineage comparisons

#### associationTest

A logical first question is whether a gene’s expression is associated with pseudotime along a given lineage, i.e., whether the smoother is flat or significantly varying along pseudotime. To address this question, the `associationTest` tests the null hypothesis that all smoother coefficients within the lineage are equal, i.e., *H*_0_: *β*_*glk*_ = *β*_*glk′*_ for all *k ≠ k′* ∈ {1,…, *K*}. This null hypothesis can be encoded in several ways; here, we chose the contrast matrix C to be an *LK × L*(*K −* 1) matrix, where each column corresponds to a contrast between two consecutive *β*_*glk*_ and *β*_*gl*(*k*+1)_ and where we have *K -* 1 contrasts per lineage for a total of *L*(*K −* 1) contrasts.

#### startVsEndTest

By default, the `startVsEndTest` compares mean expression at the progenitor state (i.e., the start of the lineage) to mean expression at the differentiated state (i.e., the end of the lineage) Wald test statistic, as described above. Specifically, C is an (*LK*) *×L* matrix, whose entry in row *k* + (*l −* 1)*K* and column *l* encodes the contrast for lineage *l* and knot *k* and is defined by *b*_*k*_ (*T*_*l,max*_ − *b*_*k*_ *T*_*l,min*_) where 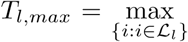 *T*_*li*_ and 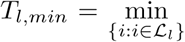 *T*_*li*_ denote, respectively, the maximum and minimum pseudotime across all cells assigned to lineage *l*. Other entries of C are set to zero. Therefore, the *l*^*th*^ element of the vector C^*T*^ *β*_*g*_ is 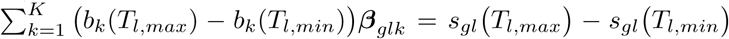, which contrasts mean expression at the beginning and at the end of the lineage. Note that contrasting the start and endpoints of a lineage is a special case of a more general capability of `tradeSeq` to compare the average expression between any two regions of a given lineage. As such, this test can be considered a generalization of cluster-based discrete DE within a lineage (e.g., Risso et al. [2018b]).

### Between-lineage comparisons

#### diffEndTest

The `diffEndTest` compares average expression at the differentiated states of multiple lineages, i.e., it compares the endpoints of different lineage-specific smoothers. It can be viewed as an analog of discrete DE for the differentiated cell types. The test is implemented using a Wald test statistic, as described above, where C is an (*LK*) × *L*(*L −* 1)*/*2 matrix. Each column of C encodes a pairwise contrast between the endpoints of two lineages, such that the corresponding element of C^*T*^ *β*_*g*_ is 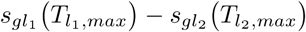 for lineages *l*_1_ and *l*_2_.

#### patternTest

This test compares the expression patterns along pseudotime between lineages by contrasting a fixed set of equally-spaced pseudotimes (*M* = 100 by default). First selecting the pseudotimes and subsequently comparing their expression levels between lineages, allows for comparisons between smoothers of different lengths. Specifically, for lineage *l*, let *P*_*lm*_ denote the *m*^th^ equally-spaced pseudotime between *T*_*l*,min_ and *T*_*l*,max_. The contrast of *M* points corresponds to testing the null hypothesis that a gene has the same expression pattern along pseudotime across the lineages under comparison, while normalizing for the length of the lineages. The test is implemented using a Wald test statistic, as described above, where C is an (*LK*) × *L*(*L −* 1)*M/*2 matrix. Each column of C encodes a pairwise comparison between two pseudotimes of two different lineages, such that the corresponding element of C^*T*^ *β*_*g*_ is 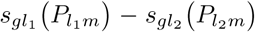 for lineages *l*_1_ and *l*_2_ and *m* ∈ {1,…, *M*}. The test is implemented through the eigendecomposition of the estimated variance-covariance matrix of the contrasts to avoid singularity problems [Smyth, 2004] (see Supplementary Methods). It should be noted that this test is a global test, able to identify both differences in patterns of expression as well as genes with similar patterns but different mean expression across the pseudotime range. It is therefore most useful as a screening test to identify any form of differential expression between the lineages.

#### earlyDETest

The `earlyDETest` aims to identify genes that are driving the differentiation around the branching. It is similar to the `patternTest`, in that it also compares the expression patterns along pseudotime between lineages by contrasting a fixed set of equally-spaced pseudotimes (*M* = 100 by default). However, instead of using points distributed from the beginning *T*_*l*,min_ to the end *T*_*l*,max_ of the lineages as in the `patternTest`, it relies on points over a shorter range of time. In the current implementation, this range is delimited by the pseudotimes of two user-defined knots. The knots should be chosen to span the branching event (or any event of interest) and do not need to be consecutive.

### Global testing

While the statistical tests introduced above can assess DE within one lineage or between a pair of lineages, one may want to investigate multiple (i.e., more than two) lineages. For example, if a trajectory consists of three lineages, one may wish to test the global null hypothesis that, for each of the three lineages, there is no association between gene expression and pseudotime using the `associationTest`. The null hypothesis that would be tested can be expressed as *H*_0_:∀*l* and ∀*k* ≠ *∀ k′, β*_*glk*_ = *β*_*glk′*_, i.e., within each of the three lineages, all *K* regression coefficients are equal. We refer to such a test as a “global test”. The `tradeSeq` package provides functionality for global testing for each of the within and between-lineage tests described above. For within-lineage tests, the user can specify whether the test should be done for each lineage individually or at the global level (i.e., for all lineages). For between-lineage tests, the user can specify whether only a pair of lineages should be assessed for DE or all pairwise comparisons should be performed.

### Stage-wise testing

For the olfactory epithelium case study [Fletcher et al., 2017] detailed below, we apply stage-wise testing, as implemented in `stageR` [Heller et al., 2009, Van den Berge et al., 2017], to assess DE between lineages using multiple tests for each gene. Stage-wise testing aims to control the overall FDR (OFDR) [Heller et al., 2009], i.e., the expected proportion of genes with at least one falsely rejected null hypothesis among all genes declared DE. In our case, the OFDR can be interpreted as a gene-level FDR [Van den Berge et al., 2017]. Stage-wise testing is performed in two stages, a screening and a confirmation stage. At the screening stage, each gene is screened by performing a global test across all null hypotheses of interest, essentially testing whether at least one of these hypotheses can be rejected. At that stage, the FDR is controlled across genes at level *α*_*I*_. At the confirmation stage, each specific hypothesis is assessed, but only for the genes that have passed the screening stage. For each gene, the family-wise error rate (FWER) is controlled across hypotheses at level 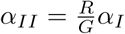, where *R* denotes the number of genes that had their global null hypothesis rejected at the screening stage and *G* the total number of genes assessed. Heller et al. [2009] proved that this procedure controls the overall FDR at level *α*_*I*_. It should be noted that, while the stage-wise testing paradigm theoretically controls the OFDR (given underlying assumptions are satisfied), the resulting *p*-values might still be too liberal since the same data are used for trajectory inference and differential expression. As mentioned before, we use *p*-values simply as numerical summaries for ranking the genes for further inspection.

### Clustering gene expression patterns

The NB-GAM can also be used to cluster genes according to their expression patterns, as shown in Figure 1. Specifically, for each gene, we extract a number of fitted values for each lineage (100 by default). We then use resampling-based sequential ensemble clustering, as implemented in`RSEC` [Risso et al., 2018b], to perform the clustering based on the first ten principal components of the standardized fitted values matrix (i.e., the fitted values are standardized to have zero mean and unit variance across cells for each gene). This clustering approach is implemented in the `tradeSeq` package (`clusterExpressionPatterns` function) for downstream analysis facilitating the interpretation of DE genes.

### Implementation

The above DE tests are implemented in the open-source R package `tradeSeq`, available on our GitHub repository (https://github.com/statOmics/tradeSeq) and to be submitted to the Bioconductor Project (http://www.bioconductor.org). We provide an extensive vignette along with the package, as well as a cheat sheet describing the different types of DE patterns detected with each test.

### Methods comparison

`slingshot` is a fast and robust method for TI that was shown to be among the top performing methods in a recent large-scale benchmarking study [Saelens et al., 2019]. Hence, we evaluate `tradeSeq` down-stream of a `slingshot` analysis, which can work with any dimensionality reduction and clustering methods. `slingshot` builds a cluster-based minimum spanning tree (MST) to infer the global lineage topology and make an initial assignment of cells to lineages. This structure is then smoothed by fitting simultaneous principal curves, which refine the assignments of cells to lineages. This process results in lineage-specific pseudotimes and weights of assignment for each cell.

`GPfates` [Lönnberg et al., 2017] is a Python package that adopts Gaussian processes in reduced dimension to infer trajectories. Dimensionality reduction is performed using Gaussian process latent variable models (GPLVM) [Lawrence, 2003]. `GPfates` is able to identify bifurcation points and assess how well a bifurcation fits the expression pattern for every gene, i.e., whether the patterns of gene expression are different between the lineages. This allows us to compare a `slingshot` + `tradeSeq` analysis with a `Gpfates` analysis. In addition, we also evaluate a `tradeSeq` analysis downstream of TI with `GPfates`, since `Gpfates` also calculates posterior probabilities that each cell belongs to a particular lineage. We then compare the complete `GPfates` (TI and DE) analysis to a `GPfates` + `tradeSeq` analysis.

`Monocle 2` [Qiu et al., 2017] applies reverse graph embedding to infer trajectories and yields a principal graph that is allowed to branch. It provides a similar approach as `tradeSeq` with the branch expression analysis modeling (`BEAM`) method. It assumes a gene-wise negative binomial model for gene expression, where the mean is expressed in terms of lineage-dependent smooth functions of pseudotime, i.e.,

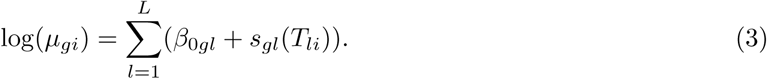

In this model, the lineage-specific intercepts *β*_0*gl*_ account for mean differences in expression between lineages, while the lineage-specific smoothers *s*_*gl*_(*t*) model the expression change along pseudotime. To test for lineage-dependent expression, the full model is compared to a null model of the form

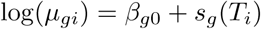

using a likelihood ratio test. Thus, `BEAM` tests whether the smooth functions of gene expression along pseudotime are different between lineages. Importantly, the `BEAM` method does not allow the inclusion of other covariates such as batch, and it is restricted to the dimensionality reduction methods that are implemented in the software package. Additionally, it only provides a screening test (like the `patternTest` in `tradeSeq`), as it only allows testing for any difference in the expression profiles between lineages and does not specify the exact type of divergence.

An alpha release for `Monocle 3` is available online (downloaded August 30, 2018 from the Monocle GitHub repository) which, unlike `Monocle 2`, performs uniform manifold approximation and projection [McInnes et al., 2018] dimensionality reduction upstream of the trajectory inference. Additionally, `Monocle 3` implements the Moran’s I test to discover genes whose expression is significantly associated with pseudotime; a functionality that is unavailable in `Monocle 2`.

`edgeR` [McCarthy et al., 2012] is a discrete differential expression method, where the groups under comparison must be defined *a priori*. It is therefore useful for assessing DE between, for example, annotated clusters or different treatment groups. For such comparisons, `edgeR` is a powerful method with high sensitivity. However, it is limited to discrete DE and cannot be applied when interested in continuous DE, e.g., assessing differences in expression patterns along pseudotime.

### Simulation study

The simulation study evaluates methods that (differentially) associate gene expression with pseudotime for three different trajectory topologies, i.e., a cyclic, a bifurcating, and a multifurcating trajectory. As independent evaluation, we use the extensive simulation framework that previously served for benchmarking trajectory inference methods in Saelens et al. [2019]. Interested readers should refer to the original publication for details on the data simulation procedure. Dataset characteristics are listed in Table 1.

**Table 1:** *Overview of simulated datasets*. Each dataset is simulated using a framework from the `dynverse` toolbox and is characterized by the topology of the trajectory, as well as the number of cells and genes. Low-dimensional representations of representative datasets can be found in Figure 2. Note that the cyclic datasets have some variation in the numbers of genes and cells and in the amount of differential expression, which is inherent to the `dyngen` simulation framework.

|  | Cyclic dataset | Bifurcating dataset | Multifurcating dataset |
| --- | --- | --- | --- |
| Simulation framework | `dyngen` | `dyntoy` | `dyntoy` |
| Number of cells | 505 – 508 | 500 | 750 |
| Number of genes | 312 – 444 | 5,000 | 5,000 |
| % of DE genes | 42 – 47% | 20% | 20% |
| Number of lineages | 1 | 2 | 3 |
| Topology | Cyclic | Bifurcating | Multifurcating |
| Number of datasets | 10 | 10 | 1 |

For each of the cyclic and bifurcating topologies, we generate and analyze 10 datasets. Since the multifurcating topology is very variable across simulations due to its flexible definition, its analysis requires substantial supervision. Therefore, we analyze only one representative multifurcating dataset.

Prior to trajectory inference, the simulated counts are normalized using full-quantile normalization [Bolstad et al., 2003, Bullard et al., 2010]. For TI with `slingshot`, we apply principal component analysis (PCA) dimensionality reduction to the normalized counts and *k*-means clustering in PCA space. For the bifurcating and multifurcating trajectories, the start and end clusters of the true trajectory are provided to `slingshot` to aid it in inferring the trajectory. For the `edgeR` analysis, we assess DE between the end clusters that are also provided to `slingshot`. The `BEAM` method can only test one bifurcation point at a time. For the multifurcating dataset, we therefore assessed both branching points separately and aggregated the *p*-values using Fisher’s method [Fisher, 2006]. For the `tradeSeq` and `edgeR` analyses of the multifurcating dataset, we perform global tests across all three lineages.

We assess performance based on scatterplots of the true positive rate (TPR) vs. the false discovery proportion (FDP), according to the following definitions

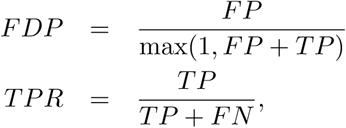

where *FN*, *FP*, and *TP* denote, respectively, the numbers of false negatives, false positives, and true positives. FDP-TPR curves are calculated and plotted with the Bioconductor R package `iCOBRA` [Soneson and Robinson, 2016].

### Case studies

#### Mouse bone marrow dataset

We use as first case study the mouse haematopoiesis scRNA-seq dataset of Paul et al. [2015]. Two small cell clusters corresponding to the dendritic and eosinophyl cell types were removed from the trajectory inference and downstream DE analysis, since these are outlying cell types that do not seem to belong to any particular lineage (Supplementary Figure S2).

`tradeSeq` downstream of `slingshot` is compared to the `BEAM` approach from `Monocle 2`. Since `BEAM` is restricted to the dimensionality reduction methods implemented in the package, we use independent components analysis (ICA) for both `slingshot` and `Monocle 2` in this comparison. For `Monocle 2`, we specify the argument `num paths=2` to aid it in inferring two lineages.

Subsequently, we demonstrate a `tradeSeq` analysis downstream of `slingshot` by performing dimensionality reduction using UMAP [McInnes et al., 2018], following the data processing pipeline described in the Monocle 3 vignette, since this better reflects the biology of the experiment.

In this case study, we show how one can perform multiple tests to identify genes with distinct types of behavior, specifically, genes that are deemed DE for one test (test 1) but not another (test 2). Let 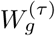 denote the test statistic for gene *g* in test *τ* ∈ {1, 2} and 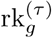 denote the rank (in terms of ordering from low to high) of 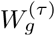 among all *G* test statistics associated with the *G* genes. Then, define a score for each gene *g* as 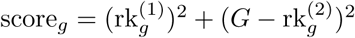. Genes with high scores are genes which are expected to be DE for test 1 but not DE for test 2 and vice versa. This is used to identify genes that are DE with the `patternTest` (test 1) but not the `diffEndTest` (test 2), i.e., genes that are transiently DE between lineages. Note that the procedure only provides a ranking of the genes and not an evaluation of statistical significance.

#### Mouse olfactory epithelium dataset

The olfactory epithelium (OE) dataset from Fletcher et al. [2017] is our second case study. We use the lineages discovered in the original manuscript. In brief, counts are normalized using full-quantile normalization [Bolstad et al., 2003, Bullard et al., 2010] followed by regression-based adjustment for quality control variables [Fletcher et al., 2017]. Dimensionality reduction is performed through PCA on the normalized log-transformed counts that are offset by 1 to avoid taking the log of zero, i.e., log(*y* + 1). Clustering is performed through *k*-means on the first 50 principal components by varying the number of clusters *k* ∈{4,…, 15}; stable clusters are derived using `clusterExperiment` [Risso et al., 2018b], yielding a final repertoire of 13 cell clusters. Next, `slingshot` is used to infer trajectories with the initial cluster chosen by known marker genes of horizontal basal cells (HBC), an adult stem cell population. A double bifurcation is discovered, with the first giving rise to sustentacular cells and two more lineages that split into microvillous cells and olfactory sensory neurons. The data were downloaded from GEO with accession number GSE95601.

## Results

In this section, we first evaluate `tradeSeq` on several simulated datasets with trajectories that span different topologies. Next, we demonstrate how `tradeSeq` can improve biological interpretation of trajectory inference results by applying it to two real datasets, a scRNA-seq dataset for mouse bone marrow [Paul et al., 2015] and a SMART-Seq2 dataset for the mouse olfactory epithelium [Fletcher et al., 2017].

### Simulation study

To benchmark relevant differential expression methods, we generated multiple datasets, spanning three distinct trajectory topologies, using the independently developed `dynverse` toolbox [Saelens et al., 2019]. We first generated 10 datasets (see Figure 2a for a representative dataset) corresponding to a single lineage contributing to a cyclic trajectory. Next, we considered a bifurcating topology, where a common lineage bifurcates into two differentiating lineages, and likewise generated 10 datasets (see Figure 2b for a representative dataset). Finally, we considered a multifurcating topology (Figure 2c). Note that the simulated datasets are relatively “clean”, as reflected by the high sensitivity and specificity of most methods. In particular, cells are approximately uniformly distributed along each lineage and balanced between lineages. The datasets are, however, still useful to provide a relative ranking of the methods.

**Figure 2:**
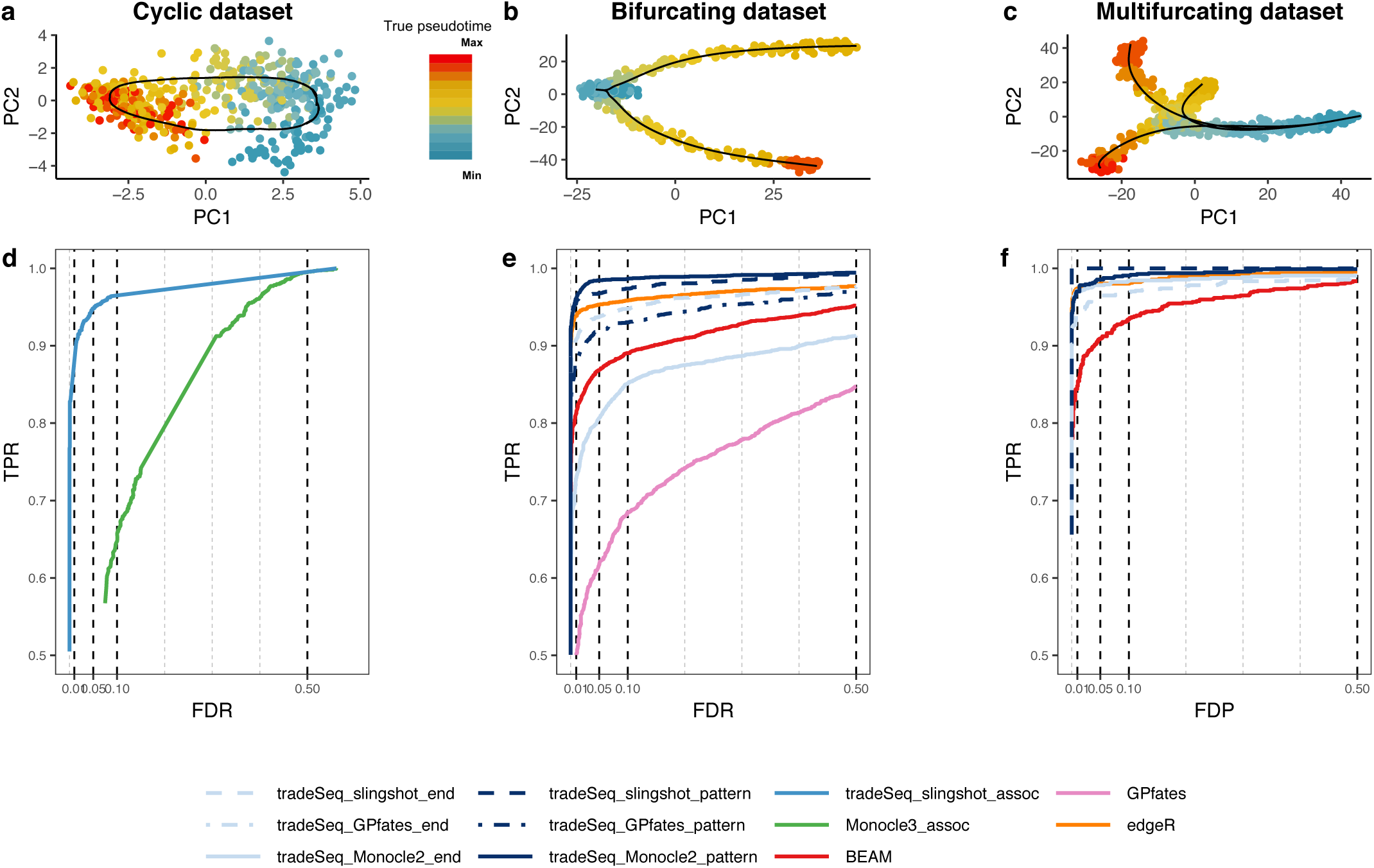
Simulation study results. PCA plots for the (a) cyclic, (b) bifurcating, and (c) multifurcating simulated trajectories. The plotting symbol for each cell is colored according to its true pseudotime; trajectories (in black) were inferred by `princurve` in (a) and `slingshot` in (b) and (c). (d-f) Scatterplot of the true positive rate (TPR) vs. the false discovery rate (FDR) or false discovery proportion (FDP) for various DE methods applied to the simulated datasets. Panel (d) displays the average performance curves of DE methods across all 10 cyclic datasets. The `associationTest` from `tradeSeq` has superior performance for discovering genes whose expression is associated with pseudotime, as compared to `Mon-ocle 3`. When investigating differential expression between lineages of a trajectory, the `patternTest` of `tradeSeq` consistently outperforms the `diffEndTest` across all three TI methods, since it is capable of comparing expression across entire lineages. Panel (e) displays the average performance curves across the three bifurcating datasets where all TI methods recovered the correct topology. Here, all `tradeSeq` `patternTest` workflows, `tradeSeq slingshot end`, and `edgeR` have similar performance and all are superior to `BEAM` and `GPfates`. Note that the performance of `tradeSeq Monocle2 end` deteriorates as compared to `tradeSeq slingshot end`; the curve for `tradeSeq GPfates end` is not visible in this panel due to its low performance. For the multifurcating dataset of panel (f), `tradeSeq slingshot` has the highest performance, closely followed by `tradeSeq Monocle2` and `edgeR`.

We demonstrate the versatility of `tradeSeq` by using it downstream of three trajectory inference methods, `slingshot` [Street et al., 2018], `Monocle 2` [Qiu et al., 2017], and `GPfates` [Lönnberg et al., 2017], which will be denoted by `tradeSeq slingshot`, `tradeSeq Monocle2`, and `tradeSeq Gpfates`, respectively. However, we find that `GPfates` fails to recover the expected trajectory topology if run in an unsupervised way (Supplementary Figure S3). Feeding the true pseudotimes as input to `GPfates` may, however, result in meaningful trajectories (Supplementary Figure S3). We therefore adopt this approach in the simulation study, but note that this may provide an *a priori* competitive advantage to `GPfates` over other TI methods and that this would be impossible for real datasets.

#### Within-lineage DE

First, we look for genes whose expression is associated with pseudotime for datasets with a cyclic topology (e.g., Figure 2a). We compare the `associationTest` of a `tradeSeq slingshot` analysis to the Moran’s I test implemented in `Monocle 3`. We only consider `Monocle 3` because it is the only method that provides a test to assess the association between gene expression and pseudotime within a single lineage. For each TI method, we use the default/recommended dimensionality reduction method, which is PCA for `slingshot` and UMAP for `Monocle 3`. `Monocle 3`, however, often fails to reconstruct the cyclic topology and instead may fit a disconnected or branching trajectory (Supplementary Figure S4). The Moran’s I test still has reasonably high sensitivity, while `tradeSeq` downstream of `slingshot` provides superior performance to discover genes whose expression is associated with pseudotime (Figure 2d).

#### Between-lineage DE

For the bifurcating datasets (Figure 2b), we assess differential expression between lineages using the `diffEndTest` and the `patternTest` from `tradeSeq`, downstream of TI methods `slingshot`, `Monocle 2`, and `GPfates`. We compare these tests with available approaches for trajectory-based differential expression analysis, namely, `BEAM` (implemented in `Monocle 2`) and `GPfates`. Furthermore, we compare against the discrete DE method `edgeR`, where we supervise the test to assess DE between the clusters at the true endpoints of each lineage, as derived through *k*-means clustering in PCA space. For each TI method, we use the default/recommended dimensionality reduction, which is PCA for `slingshot`, GPLVM for `GPfates`, and DDRTree for `Monocle 2`.

`Monocle 2` and `GPfates` fail to detect the correct topology of the trajectory (i.e., a bifurcation) in, respectively, 3 and 4 out of the 10 datasets (Supplementary Figures S6 and S7). In addition, out of the remaining 7 datasets, `Monocle 2` misplaces the bifurcation in 4 of them, causing the two simulated lineages to be merged into the same lineage and creating another incorrect lineage. This strongly obscures the DE testing results (Supplementary Figure S6). `slingshot`, on the other hand, correctly identifies the topology and reconstructs the trajectory for all 10 datasets.

Figure 2e shows performance curves for the three datasets for which all methods are able to recover the true topology of the simulated trajectory. The `tradeSeq` `patternTest` has superior performance regardless of the TI method. Only `edgeR` achieves a similar performance. This is not surprising since the `edgeR` analysis is supervised to compare the true cell populations at the endpoints of the lineages. Interestingly, `tradeSeq`’s `diffEndTest` based on the `slingshot` trajectory performs comparably to a supervised `edgeR` analysis. This is especially encouraging, since the `diffEndTest` is a smoother-based analog of discrete DE. For TI methods `Monocle 2` and `GPfates`, `diffEndTest` performs poorly, which is not surprising since the endpoints are typically ill-defined or artificially extended in the estimated trajectories for those methods (Supplementary Figures S6 and S7). In general, `BEAM` and `GPfates` are outperformed by the other methods. Across all methods, `tradeSeq slingshot` has the best performance. Finally, we recapitulate that the performance curves in Figure 2e do not provide a complete view of method performance, since 7 out of 10 datasets were not used because at least one method failed to recover the simulated trajectory. Supplementary Figure S8 shows mean performance curves across all 10 datasets for all methods, which clearly demonstrates the superiority of `tradeSeq` as a DE method and of `slingshot` as an upstream TI method. The performance and trajectories for all 10 individual datasets are shown in Supplementary Figure S9.

For the multifurcating dataset, we forego a comparison with `GPfates`, since it is restricted to discovering only a single bifurcation (Supplementary Figure S10). The `patternTest` from `tradeSeq slingshot` and `tradeSeq Monocle2` have highest performance, closely followed by `edgeR` and the `diffEndTest` for those respective TI methods (Figure 2f). `BEAM` was found to have the lowest performance.

Taken together, these results suggest that `tradeSeq` is a powerful and flexible procedure for assessing DE along and between lineages. Although `tradeSeq` is modular and can be used downstream of any TI method that provides pseudotime estimates, the choice of dimensionality reduction and TI method is crucial for the performance of the downstream analysis. The best performance was found for a `trade-Seq slingshot` analysis, so for the real datasets we will mainly focus on `slingshot` as TI method.

### Case studies

#### Mouse bone marrow dataset

Paul et al. [2015] study the evolution of gene expression for myeloid progenitors in mouse bone marrow. They construct a reference compendium of marker genes that are indicative of development from myeloid multipotent progenitors to erythrocytes and several types of leukocytes.

In order to compare our approach with `BEAM`, we are restricted to the dimensionality reduction procedures implemented in `Monocle 2`. We will therefore first apply ICA as dimensionality reduction method (Figure 3a) in the ‘Discovering cell type markers’ paragraph. Note that this approach does not fully preserve the underlying biology. Indeed, a 2D visualization of the ICA dimensionality reduction shows that there is a seemingly large gap between the multipotent progenitors and the remaining cell types, and a number of erythrocytes and granulocyte-macrophage progenitors (GMP) are misclassified as multipotent progenitors. In addition, megakaryocytes, which are thrombocyte progenitors and as such should not belong to any of the two lineages, seem to be split between the erythrocyte and leukocyte lineages. However, when applying UMAP dimensionality reduction (Figure 3b), these issues seem to be resolved and the trajectory seems to be better represented. In subsequent sections, we will therefore demonstrate the powerful interpretation of a `tradeSeq slingshot` analysis based on UMAP dimensionality reduction. This additionally demonstrates the flexibility of `tradeSeq` (and `slingshot`) to be applied downstream of any dimensionality reduction method.

**Figure 3:**
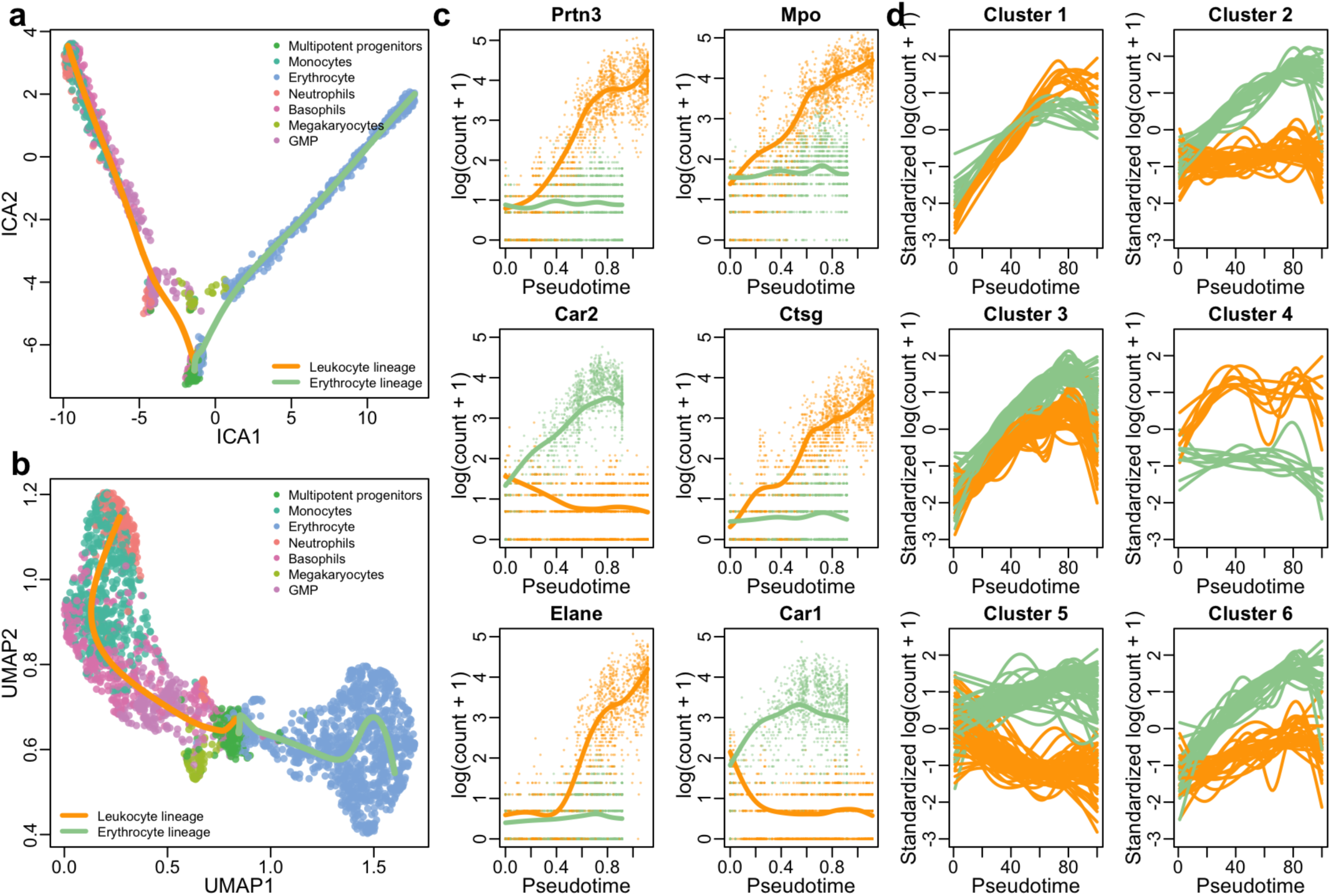
Mouse bone marrow case study [Paul et al., 2015]. (a) Two-dimensional representation of a subset of the data using independent components analysis (ICA). The myeloid trajectory inferred by `slingshot` is displayed. (b) Two-dimensional representation of a subset of the data using UMAP. The myeloid trajectory inferred by `slingshot` is displayed. The UMAP dimensionality reduction method better captures the smooth differentiation process than ICA. (c) Estimated smoothers for the top six genes identified by the `tradeSeq` `patternTest` procedure on the trajectory from (b). (d) Six clusters for the top 500 genes with different expression patterns between the two lineages (as identified by `patternTest` from `tradeSeq`).

#### Discovering cell type markers

`tradeSeq` provides the flexibility to test several interesting and distinct hypotheses for this dataset, that cannot always be considered with other methods. For instance, we can find marker genes for the progenitor cell population vs. the differentiated leukocytes or erythrocytes with the `startVSEndTest` procedure (results shown in Supplementary Figure S11). We can also discover marker genes for the differentiated cell types by comparing the differentiated leukocyte and erythrocyte cells themselves through contrasting the endpoints of the smoothers with the `diffEndTest` procedure. For the latter, `tradeSeq` finds 2,543 significantly differentially expressed genes at a 5% nominal FDR level, while `BEAM` discovers 584 genes at a 5% nominal FDR level when testing whether the association between gene expression and pseudotime depends on the lineage (Benjamini and Hochberg [1995] FDR-controlling procedure).

Since the identification of a larger set of DE genes does not necessarily imply more relevant biology, we select carefully constructed gene sets from deGraaf et al. [2016] to perform gene set enrichment analysis (GSEA) on blood cell types. As we are comparing erythrocytes with a mixture of leukocytes, we expect gene sets related to erythrocytes to be significant. Indeed, the erythrocyte gene set is the only one to be found significant by `fgsea` [Sergushichev, 2016] for the `tradeSeq` analysis (FDR adjusted *p*-value *<* .001, with normalized enrichment score of 1.59), while no significant gene sets are found for the `BEAM` analysis (as reference, in that case, the FDR adjusted *p*-value for the erythrocyte gene set is 0.56). In this case, `tradeSeq` is therefore better able to recover a meaningful biological signal.

If one assumes that the cell type labels are known for all cells in the dataset, a cluster-based comparison is possible, where the different clusters correspond to the identified cell types. We use `edgeR` [McCarthy et al., 2012] to assess differential expression between erythrocytes and neutrophils, since this comparison is most analogous to `tradeSeq`’s `diffEndTest`. Only `edgeR` finds evidence for gene sets related to eosinophils and T-cells (FDR adjusted *p*-values of 0.043 and 0.049, respectively), however, the eosinophil cells were removed from this dataset prior to analysis (see Methods, subsection ‘Case studies: Mouse bone marrow dataset’). The GSEA results for `edgeR` also provide less evidence for erythrocytes (FDR adjusted *p*-value = 0.043, normalized enrichment score=1.21) as compared to the `tradeSeq` analysis. None of the methods, however, recover evidence for the neutrophil cell types that are identified at the end of the lineage (Figure 3b; FDR adjusted *p*-values *p*_tradeSeq_ = 0.54, *p*_BEAM_ = 0.85, and *p*_edgeR_ = 0.81). A `tradeSeq` analysis can thus provide relevant biological results without using the cell type labels. Moreover, while a cluster-based comparison can be powerful in some cases, many hypotheses are difficult to assess with discrete DE, as we demonstrate in the following paragraphs.

#### Discovering progenitor population markers

In addition to looking for markers at the differentiated cell type level, we could also look for markers of developing myeloid cells. `tradeSeq` can accommodate this by identifying genes with significantly different expression patterns between lineages. Remarkably, the top six genes (*Prtn3, Mpo, Car2, Ctsg, Elane*, and *Car1*, Figure 3b) are all confirmed as biomarkers in the extensive analysis of the original manuscript of Paul et al. [2015], confirming the relevant ranking of `patternTest` in `tradeSeq`. Indeed, *Prtn3* was found to be monocyte-specific, while *Mpo* and *Car2* discriminated between erythroid lineage progenitors and myeloid lineage progenitors. The cluster of genes *Elane, Prtn3*, and *Mpo* were the strongest markers for myeloid lineage progenitors and monocytes. Summarized, all six top genes were labelled as “key genes” for hematopoiesis [Paul et al., 2015].

It might also be interesting to examine genes with significantly different expression patterns, that show little evidence for DE at the endpoints. We therefore select genes with both a high Wald test statistic (low *p*-value) for the `patternTest` and a low test statistic (high *p*-value) for the `diffEndTest`. Following the approach described in ‘Case studies: Mouse bone marrow dataset’ (Methods), we assign a score for each gene. Remarkably, the five genes with the highest scores (*Irf8, ApoE, Erp29, Psap*, and *Lamp1*) were previously found to be major regulators of hematopoiesis. Indeed, *Irf8* has previously been identified as a major transcription factor involved in myeloid lineage commitment [Kurotaki et al., 2014, Paul et al., 2015]. *Erp29, Psap*, and *Lamp1* are all direct targets of the *Irf8* transcription factor [Paul et al., 2015, Shin et al., 2011, Marquis et al., 2011], while *ApoE* regulates stem cell proliferation in atherosclerotic mice [Murphy et al., 2011] and was also identified as a marker gene in the original manuscript of Paul et al. [2015]. Note that this analysis is not possible with any other method available, since these only test for global differential gene expression between lineages.

#### Gene expression families

Modeling gene expression in terms of smooth functions of pseudotime opens the door for additional downstream interpretation of results that are impossible with discrete DE methods. For example, we can cluster the expression patterns for all genes that were deemed significant by `tradeSeq`’s `patternTest` (see Methods, section ‘Clustering gene expression patterns’). This identifies gene families that have similar expression patterns within every lineage, and also similar fold-changes between the two lineages (Figure 3c shows six random clusters). These gene sets can then be further screened for interesting patterns and validated by the biologist. Note that, for instance, the expression smoothers can be used to assess specific transient changes in expression during development, the signal for which might be diluted in cluster-based DE.

#### Mouse olfactory epithelium dataset

Fletcher et al. [2017] study the development of horizontal basal cells (HBC) in the olfactory epithelium (OE) of mice. They activate the HBCs to be primed for development, which subsequently give rise to three different cell types: sustentacular cells, microvillous cells, and olfactory sensory neurons. The olfactory sensory neurons are connected to the olfactory bulb for signal transduction of smell and the sustentacular cells are general supportive cells in the OE. The function of microvillous cells, however, is not well understood; while some cells have axons ranging to the olfactory bulb, potentially indicating a sensory neuron function, others lack a basal process or axon [Hegg et al., 2010]. The samples from Fletcher et al. [2017] were processed using the Fluidigm C1 system with SMART-Seq library preparation, hence we expect zero inflation to be present in this dataset. We therefore fit ZINB-GAMs to analyze the data using `tradeSeq` downstream of `slingshot`. Zero inflation weights are estimated with the ZINB-WaVE method [Risso et al., 2018a], using the cluster labels and batch as covariates. Note that no other trajectory-based DE method can account for zero inflation or provide the range of tests available in `tradeSeq`; hence, we forgo a comparison with other methods aside from a ZINB-edgeR analysis [Van den Berge et al., 2018].

#### Within-lineage DE

We first consider differential expression along the neuronal lineage (the orange lineage in Figure 4a). Using the `associationTest` implemented in `tradeSeq`, we recover 1,951 genes at a 5% nominal FDR level. Within the top DE genes, clear clusters of expression can be observed (Figure 4c), that are more active either at the beginning, at specific locations along the lineage, or at the end of the lineage. Since Fletcher et al. [2017] observed that cells associated with the neuronal lineage undergo mitotic division during differentiation, we investigate whether we can recover the cell cycle biology using the `associationTest`. Indeed, many of the top genes are related to the cell cycle (Supplementary Figure S12).

**Figure 4:**
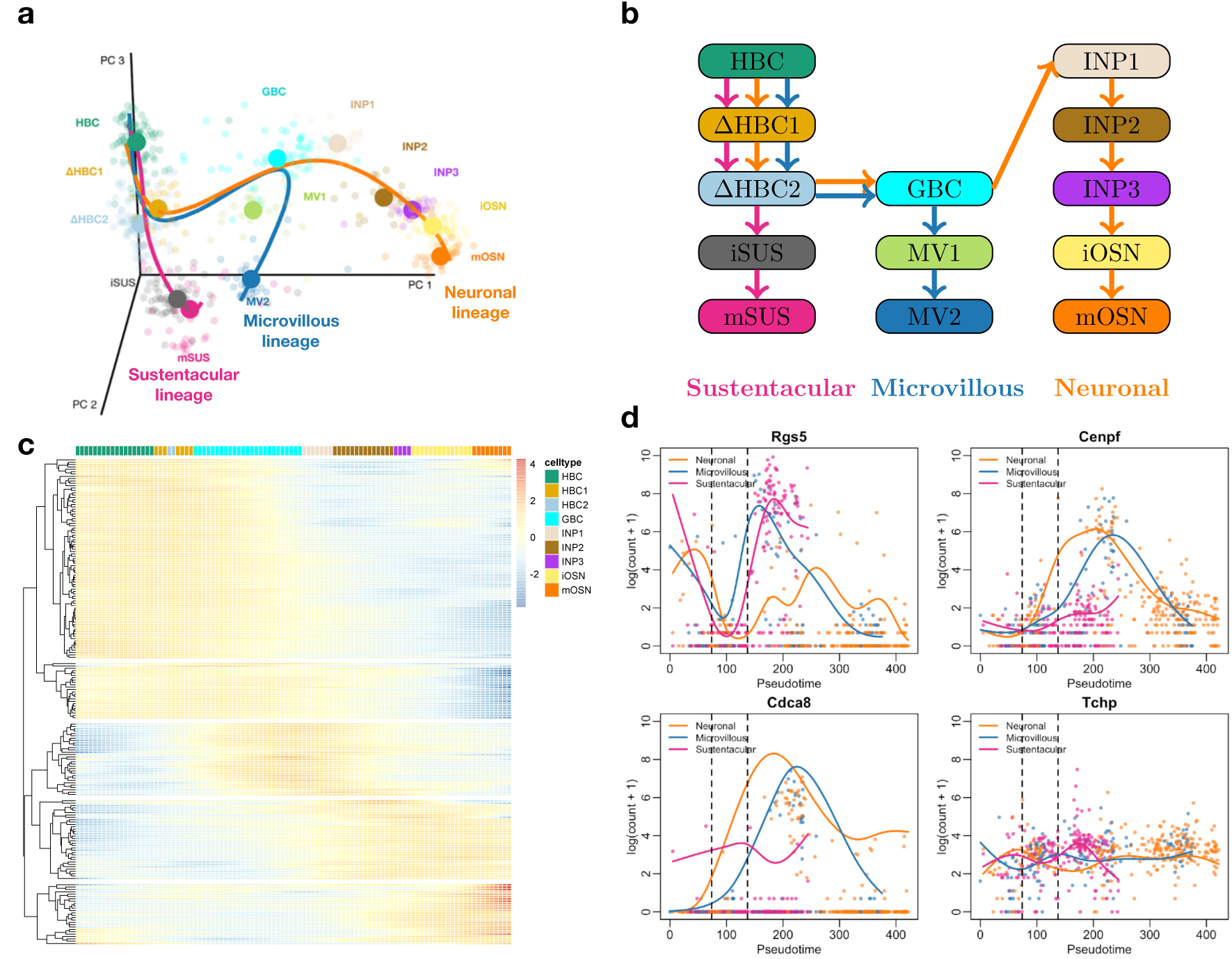
Mouse olfactory epithelium case study [Fletcher et al., 2017]. (a) Three-dimensional PCA plot of the scRNA-seq data, where cells are colored according to their cluster membership as defined in the original paper (see Methods). The simultaneous principal curves for the lineages inferred by `slingshot` are displayed. (b) Schematic of the cell types and their ordering along the lineages. (c) Heatmap for the top 200 genes that are associated with the neuronal lineage, as identified with the `associationTest` procedure from `tradeSeq`. Five clear gene clusters can be identified, each with a different region of activity during the developmental process. (d) Four out of a total of 16 genes that are significant in all pairwise comparisons with `earlyDETest` between the pseudotimes of knots 2 and 4 (knots indicated with vertical dashed lines). All 16 genes are plotted in Supplementary Figure S14.

We also seek biological markers that differentiate the progenitor cells from the differentiated cell types in any of the three lineages using the `startVsEndTest` procedure as part of a global test (see Methods) (i.e., gene expression is compared between the start and end states for each lineage and the evidence is aggregated across the three lineages using a global test) and then perform GSEA on the top 250 genes. The results for the top 20 gene sets (Supplementary Table S1) clearly reflect the biology of the experiment. The HBCs were primed for differentiation and the top three gene sets correspond to responses to (organic, external, and endogenous) stimuli. In addition, neurogenesis and tissue development are the fourth and fifth most significant gene sets, respectively, while the remaining list contains sets related to cell development, differentiation, and epithelium development, amongst others. Although the neuronal and microvillous cell lineages undergo mitotic division, it is worth noting that cell cycle related gene sets are absent from the `startVsEndTest` results, since this process occurs during differentiation, but not in the resting HBC or differentiated cell populations.

#### Between-lineage DE

We compare the three lineages by assessing differences in their expression patterns through stage-wise testing with the `patternTest` procedure (see ‘Case studies: Mouse olfactory epithelium dataset’ in Methods). At the screening stage, we first test whether any two lineages have significantly different expression patterns. The genes that pass the screening stage are then further assessed to discover which specific pair of lineages are deviating in their expression pattern. The screening hypothesis discovers 1,772 genes that have different expression patterns between any pair of lineages, at 5% nominal FDR level (as reference, the top six genes are plotted in Supplementary Figure S13). As could be expected, a large majority of genes (1,446) are significant in the neuronal-sustentacular lineage comparison. However, remarkably, we discover more DE genes when comparing the microvillous and neuronal lineages (891 genes) than when comparing the microvillous and sustentacular lineages (581 genes), even though the microvillous lineage shares a longer path with the neuronal lineage. Out of all significant genes, 288 genes were identified in all three pairwise comparisons. Investigating the top 20 gene sets from GSEA based on the MSigDB database reveals that 15*/*20 of the top gene sets are related to the mitotic cell cycle (Supplementary Table S2). This is reassuring, since only the neuronal and microvillous lineages undergo cell cycle, according to Fletcher et al. [2017]. In addition, we find gene sets related to neurogenesis, referring to the development of olfactory sensory neurons, and response to external stimuli, referring to the priming of HBCs for differentiation. The functional interpretation of the results from the combined ZINB and `tradeSeq` analysis hence confirms the biology of the experiment and the battery of possible tests unlock a more detailed and meaningful interpretation of the results.

None of the previously developed trajectory-based methods for assessing differential expression between lineages can accommodate zero inflation. The only relevant comparison is between the `diffEndTest` procedure from `tradeSeq` and a discrete DE test between the differentiated cell types using ZINB-`edgeR` as introduced in Van den Berge et al. [2018]. For both methods, we use a global test to compare mean expression between all three differentiated cell types. While a ZINB-`edgeR` analysis discovers 1, 925 genes, the ZINB-`tradeSeq` analysis discovers 3, 614 genes, which include ∼ 86% of the genes also discovered by the ZINB-`edgeR` analysis. In order to assess the relevance of the extra 1, 959 genes discovered with ZINB-`tradeSeq`, we perform GSEA on this gene set (Supplementary Table S3). The top 20 significant gene sets contain relevant biological processes for the system under study, such as “regulation of multicellular organismal development”, “positive regulation of biosynthetic process”, “regulation of cell differentiation”, and “tissue development”.

We can also identify genes that drive the branching based on `earlyDETest` applied around the first branching point (see Figure 4). We apply stage-wise testing [Van den Berge et al., 2017] (see ‘Case studies: Mouse olfactory epithelium dataset’ in Methods) to first assess any difference across the three lineages using a global test. We discover 804 genes to be DE between any of the three lineages at a 5% nominal FDR level. Among those 804 genes, we then discover 415 significant genes between the neuronal and microvillous lineages, 353 significant genes between the microvillous and sustentacular lineages, and 238 significant genes between the neuronal and microvillous lineages (Supplementary Figure S15). Only 16 genes are significant in all pairwise comparisons (Supplementary Figure S15), and these genes may be potentially important regulators of the transcriptional program involved in olfactory epithelium development. One gene, *Znf7* indeed is known for its DNA binding transcription factor activity. Additionally, *Gldn* promotes formation of the nodes of Ranvier in the peripheral nervous system and *Tchp* is involved in cytoskeleton remodeling of neurofilaments. *Add2* helps in the assembly of a spectrin-actin network at sites of cell-cell contact in epithelial tissues, *Rgs5* is involved in the induction of endothelial apoptosis, and *Frem1* encodes a basement membrane protein that may play a role in craniofacial development. More generally, *Fgf13* is involved in a variety of biological processes, including embryonic development, cell growth, morphogenesis; *Racgap1* plays a regulatory role in cytokinesis, cell growth, and differentiation.

## Discussion

We have proposed `tradeSeq`, a novel suite of tests for identifying dynamic temporal gene regulation using single-cell RNA-seq data. These tests allow researchers to investigate a range of hypotheses related to temporal gene expression, ranging from the general to the highly specific. Whereas previous methods only provide global tests of differential expression along or between lineages, `tradeSeq` offers a highly flexible framework that can be adapted to a single lineage, multiple lineages, or specific points or ranges along lineages. The flexibility provided by `tradeSeq` is crucial, as trajectory-based DE is often the final (or near final) step in a much longer analysis pipeline.

Our analyses are based on the NB-GAM of Equation (1) which conditions on cell pseudotimes and hence ignores the fact that pseudotimes are typically inferred random variables. We therefore expect some uncertainty in pseudotime values which may or may not be quantified by a particular TI method. Even when measures of pseudotime variability are available, neither `tradeSeq` nor other methods such as `BEAM` and `GPFates` currently make use of this information. Instead, all of these methods treat the pseudotimes as fixed and known.

While we generally assume that pseudotime values are on similar scales across lineages, this may not always be the case. Trapnell et al. [2014] noted that any trajectory inference method can produce pseudotime values that are not necessarily reflective of true biological time. At best, pseudotime values represent some monotonic transformation of the true maturity of each cell. Therefore, some authors have proposed the use of dynamic time warping to optimally align pseudotime values from different experiments on potentially different scales [Alpert et al., 2018]. This approach can be beneficial in cases where, for example, one lineage is much longer or shorter than another. If a gene, in reality, has a similar pattern of expression along two such lineages, this pattern could, for instance, consume 75% of the shorter lineage, but only 25% of the longer lineage. As such, the gene could be called DE by the `patternTest` procedure. However, applying the same test after dynamic time warping may yield a negative result. Since `tradeSeq` only requires the estimated pseudotimes as input, which could be warped or not, it is compatible with any form of warping between lineages. We urge users to carefully consider whether pseudotime values across lineages are comparable and, if not, consider such warping strategies before comparing patterns of expression with `tradeSeq`.

Moving forward, it may be possible to fit ZINB-GAMs in a single step by numerically maximizing the ZINB-GAM likelihood. This could improve upon the two-step approach that we have taken in this paper, where (i) posterior probabilities of zero inflation are first estimated using ZINB-WaVE and (ii) subsequently used to unlock the NB-GAM for DE analysis in the presence of excess zeros.

In this manuscript, we have demonstrated `tradeSeq` on several scRNA-seq datasets. However, the tests that we provide downstream of the `fitGAM` function are applicable beyond this setting. Indeed, the framework may also be applicable to, e.g., downstream analysis of chromatin accessibility trajectories in scATAC-seq datasets (e.g. Chen et al. [2019]) or bulk RNA-seq time-course studies.

While we propose a number of tests based on the NB-GAM, it is important to realize that users may also implement their own statistical tests related to their specific hypotheses of interest. For example, it may be of interest to investigate whether the speed or acceleration in expression varies significantly along or between lineages. This can be assessed in the `tradeSeq` framework using first or second derivatives of the linear predictor in Equation (1), respectively. The derivatives are linear combinations of the parameters in the basis function expansion, i.e., 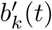, where the derivatives of the basis functions 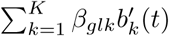 soften have a closed-form expression (e.g., cubic splines) or otherwise can be approximated using finite differencing (e.g., thin-plate splines) [Wood, 2017]. Genes that significantly increase (decrease) in their rate of expression along a lineage can then, for example, be discovered by testing whether the first derivative is significantly higher (lower) than zero. We therefore welcome contributions of new tests to the GitHub repository of the package.

Single-cell RNA-seq tends to produce noisy data requiring long analysis pipelines in order to glean biological insight. While “all-in-one” tools that simplify this analysis may be attractive from a user’s standpoint, they are not guaranteed to offer the best methods for each individual step. We therefore propose a more modular approach that expands upon previous work and opens up new classes of questions to be asked and hypotheses to be tested.

## Acknowledgments

We thank Valentine Svensson for his help on the implementation and output of the `GPfates` method. Olivier Thas and Catalina Vallejos provided constructive feedback on a draft of the manuscript. KVdB, WS, and RC are supported by the Research Foundation Flanders (no. 1S41816N, 11Z4518N, and 11Y6218N, respectively). YS is an ISAC Marylou Ingram scholar.

## Authors’ contributions

KVdB, HRdB, KS, SD, and LC conceived and designed the study. KVdB and HRdB implemented the method. KVdB, HRdB, and KS analyzed the data. WS, RC, and YS generated the synthetic data. KVdB, HRdB, KS, SD, and LC wrote the manuscript, and WS, RC, and YS contributed to revisions of an initial draft.

## Code availability

The code to reproduce the figures, tables, and analyses in the paper is available on GitHub at https://github.com/statOmics/tradeSeqPaper.

## Data availability

The code to generate all simulated datasets is included in the GitHub repository of the paper at https://github.com/statOmics/tradeSeqPaper. The data for the mouse bone marrow case study were downloaded from http://trapnell-lab.gs.washington.edu/public_share/valid_subset_GSE72857_cds2.RDS. The raw data for the olfactory epithelium case study are available on GEO with accession number GSE95601.

