## Supplementary Material for "Trajectory-based differential expression analysis for single-cell sequencing data"

### Supplementary Methods

#### Eigenvalue decomposition of $\hat{\boldsymbol{\Sigma}}_{\hat{\boldsymbol{\beta}}_g}$

Let $\mathbf{C}$ correspond to the $(LK) \times L(L-1)M/2$ matrix that defines the linear contrasts of interest for the `patternTest`, i.e., every column corresponds to the comparison of two points for a pair of lineages. All tests are implemented using a Wald test statistic defined as

$$W_g = \hat{\boldsymbol{\beta}}_g^T \mathbf{C}(\mathbf{C}^T \hat{\boldsymbol{\Sigma}}_{\hat{\boldsymbol{\beta}}_g} \mathbf{C})^{-1} \mathbf{C}^T \hat{\boldsymbol{\beta}}_g,$$

with $\hat{\boldsymbol{\Sigma}}_{\hat{\boldsymbol{\beta}}_g}$ the estimated variance-covariance matrix of the estimated smoother coefficients. Letting $\hat{\boldsymbol{\alpha}}_g = \hat{\boldsymbol{\beta}}_g^T \mathbf{C}$ and $\hat{\boldsymbol{\Sigma}}_{\hat{\boldsymbol{\alpha}}_g} = \mathbf{C}^T \hat{\boldsymbol{\Sigma}}_{\hat{\boldsymbol{\beta}}_g} \mathbf{C}$, we can rewrite the Wald statistic as

$$W_g = \hat{\boldsymbol{\alpha}}_g (\hat{\boldsymbol{\Sigma}}_{\hat{\boldsymbol{\alpha}}_g})^{-1} \hat{\boldsymbol{\alpha}}_g^T.$$

Taking the eigendecomposition of $\hat{\boldsymbol{\Sigma}}_{\hat{\boldsymbol{\alpha}}_g}$,

$$W_g = \hat{\boldsymbol{\alpha}}_g \hat{\mathbf{V}}_g^T \hat{\boldsymbol{\Lambda}}_g^{-1} \hat{\mathbf{V}}_g \hat{\boldsymbol{\alpha}}_g^T,$$

where $\hat{\mathbf{V}}_g$ is an $L(L-1)M/2 \times L(L-1)M/2$ matrix with columns corresponding to the $L(L-1)M/2$ eigenvectors of $\hat{\boldsymbol{\Sigma}}_{\hat{\boldsymbol{\alpha}}_g}$ and $\hat{\boldsymbol{\Lambda}}_g$ the $L(L-1)M/2 \times L(L-1)M/2$ diagonal matrix of eigenvalues $\lambda_i \in [0,1]$ of $\hat{\boldsymbol{\Sigma}}_{\hat{\boldsymbol{\alpha}}_g}$, in decreasing order. Note that, since $\hat{\boldsymbol{\Lambda}}_g$ is a diagonal matrix, we can simply invert its diagonal elements instead of inverting the full matrix. This provides an efficient computation of the test statistic and avoids singularity problems with the estimated variance-covariance matrix of the contrasts [Smyth, 2004]. We determine the rank $r$ of $\hat{\boldsymbol{\Sigma}}_{\hat{\boldsymbol{\alpha}}_g}$ by calculating the number of eigenvalues that are larger than $1e^{-8}\lambda_1$, with $\lambda_1$ corresponding to the largest eigenvalue. When the rank $r$ of $\hat{\boldsymbol{\Sigma}}_{\hat{\boldsymbol{\alpha}}_g}$ is less than $L(L-1)M/2$ (i.e., the matrix is not of full rank), we only use the first $r$ eigenvectors from $\hat{\mathbf{V}}_g$, associated with the largest $r$ eigenvalues from $\hat{\boldsymbol{\Lambda}}_g$.

#### Defining $\mathbf{Z}$ based on user-supplied weights

If one has user-supplied weights $\mathbf{W} = (W_{li} \in [0,1] : l \in \{1,\dots,L\}, i \in \{1,\dots,n\})$ for the assignment of cells to lineages, one can construct the binary matrix $\mathbf{Z}$ from $\mathbf{W}$ as follows.

First, note that the weights $\mathbf{W}$ may be defined differently depending on the TI method that was used to estimate them. For example, slingshot [Street et al., 2018] defines weights based on the distance from a cell to a particular lineage; hence, the sum of the weights across all lineages for a particular cell may be greater than 1. As such, these weights cannot be interpreted as probabilities. GPfates [Lönnberg et al., 2017], however, does return posterior probabilities that a cell $i$ belongs to a particular lineage $l$, where $\sum_{l=1}^{L} W_{li} = 1$ for each $i$. We therefore first normalize the weights for each cell, such that, for normalized weights $W_{li}^*$, the sum across lineages equals one, i.e., $\sum_{l=1}^{L} W_{li}^* = 1$ for each cell $i$. Next, we assign each cell $i$ to a lineage by sampling one observation from a Multinomial distribution with $L$ groups and probabilities $W_{li}^*$. The lineage assignments are then encoded in the $L \times n$ matrix $\mathbf{Z}$, by setting all elements of the $i^{\text{th}}$ column equal to zero except for a 1 in the row corresponding to the sampled lineage for cell $i$.

The multinomial sampling to assign each cell to a lineage may introduce variability in the results if the models are fit multiple times, due to differing cell allocations. This is especially so if there is a high uncertainty about the lineage allocation (e.g., a cell is equally likely to belong to each lineage), which typically occurs around the inception of a trajectory. While we ensure reproducibility by setting a seed in the software, the results may vary slightly over different seeds. To quantify this variability, we use the data of Paul et al. [2015] and allocate cells to lineages using 10 different seeds. Since we expect the variability across different assignments to be largest at the inception of the lineage, we evaluate DE using tradeSeq's `startVsEndTest`. Using a global test across the two lineages, the number of DE genes at a 5% nominal FDR level varied between 1,990 and 2,049, with 1,739 DE genes shared across all 10 assignments (Supplementary Figure S16). For each assignment, at least 93% of the 1,000 top DE genes are shared with the top 1,000 DE genes of any other assignment.

### Supplementary Tables and Figures

Table S1: *Mouse olfactory epithelium dataset.* The top 20 significant GO sets for the top 250 genes when assessing global differential expression between the progenitor and differentiated cell populations using the `tradeSeq` `startVsEndTest` procedure. The "Overlap" column records the number of genes, out of the 250 top genes, that are included in a particular gene set. The significance of a gene set is measured by a $q$-value, obtained by applying gene set enrichment analysis (GSEA, Subramanian et al. [2005]) with the Molecular Signatures Database v6.2 (`http://software.broadinstitute.org/gsea/msigdb`).

| | **Gene set** | **Overlap** | **Genes in set** | **$q$-value** |
|---|---|---|---|---|
| 1 | cellular response to organic substance | 48 | 1848 | 1.37E-18 |
| 2 | response to external stimulus | 47 | 1821 | 2.61E-18 |
| 3 | response to endogenous stimulus | 42 | 1450 | 5.05E-18 |
| 4 | neurogenesis | 41 | 1402 | 8.48E-18 |
| 5 | tissue development | 42 | 1518 | 1.65E-17 |
| 6 | regulation of cellular component movement | 31 | 771 | 6.92E-17 |
| 7 | negative regulation of response to stimulus | 39 | 1360 | 9.55E-17 |
| 8 | positive regulation of cell communication | 41 | 1532 | 1.03E-16 |
| 9 | cellular response to endogenous stimulus | 34 | 1008 | 1.52E-16 |
| 10 | regulation of multicellular organismal development | 42 | 1672 | 2.78E-16 |
| 11 | regulation of cell differentiation | 39 | 1492 | 1.42E-15 |
| 12 | negative regulation of cell communication | 35 | 1192 | 2.42E-15 |
| 13 | neuron differentiation | 30 | 874 | 9.85E-15 |
| 14 | positive regulation of gene expression | 40 | 1733 | 2.73E-14 |
| 15 | regulation of intracellular signal transduction | 39 | 1656 | 3.36E-14 |
| 16 | negative regulation of multicellular organismal process | 30 | 983 | 1.86E-13 |
| 17 | cell development | 35 | 1426 | 3.91E-13 |
| 18 | regulation of anatomical structure morphogenesis | 30 | 1021 | 4.31E-13 |
| 19 | epithelium development | 29 | 945 | 4.31E-13 |
| 20 | response to abiotic stimulus | 30 | 1024 | 4.38E-13 |

Table S2: *Mouse olfactory epithelium dataset.* The top 20 significant GO sets for the 288 genes that were found to be significant in all pairwise comparisons between the three trajectories using the `tradeSeq` `patternTest` procedure. The "Overlap" column records the number of genes, out of the 288 top genes, that are included in a particular gene set. The significance of a gene set is measured by a $q$-value, obtained by applying gene set enrichment analysis (GSEA, Subramanian et al. [2005]) with the Molecular Signatures Database v6.2 (`http://software.broadinstitute.org/gsea/msigdb`).

| | **Gene set** | **Overlap** | **Genes in set** | $q$**-value** |
|---|---|---|---|---|
| 1 | cell cycle | 52 | 1316 | 6.8E-27 |
| 2 | cell cycle process | 48 | 1081 | 6.8E-27 |
| 3 | mitotic cell cycle | 38 | 766 | 1.66E-22 |
| 4 | organelle fission | 26 | 496 | 2.34E-15 |
| 5 | regulation of cell cycle | 33 | 949 | 8.9E-15 |
| 6 | mitotic nuclear division | 22 | 361 | 3.11E-14 |
| 7 | cell division | 24 | 460 | 3.21E-14 |
| 8 | positive regulation of molecular function | 41 | 1791 | 9.67E-13 |
| 9 | chromosome segregation | 18 | 272 | 3.9E-12 |
| 10 | sister chromatid segregation | 15 | 176 | 1.58E-11 |
| 11 | microtubule based process | 22 | 522 | 3.33E-11 |
| 12 | cellular response to stress | 36 | 1565 | 3.52E-11 |
| 13 | nuclear chromosome segregation | 16 | 228 | 3.52E-11 |
| 14 | chromosome organization | 29 | 1009 | 4.35E-11 |
| 15 | negative regulation of cell cycle | 20 | 433 | 6.38E-11 |
| 16 | regulation of cell cycle process | 22 | 558 | 8.66E-11 |
| 17 | positive regulation of gene expression | 37 | 1733 | 1.04E-10 |
| 18 | cell cycle phase transition | 16 | 255 | 1.43E-10 |
| 19 | neurogenesis | 33 | 1402 | 1.46E-10 |
| 20 | response to external stimulus | 37 | 1821 | 3.79E-10 |

Table S3: *Mouse olfactory epithelium dataset.* The top 20 significant GO sets based on the unique 1,959 genes that were only discovered with the ZINB-`tradeSeq` analysis, and not the ZINB-`edgeR` analysis, when comparing mean expression between the end points of the lineages using the `tradeSeq` `diffEndTest` procedure. The "Overlap" column records the number of genes, out of the 1,959 top genes, that are included in a particular gene set. The significance of a gene set is measured by a $q$-value, obtained by applying gene set enrichment analysis (GSEA, Subramanian et al. [2005]) with the Molecular Signatures Database v6.2 (`http://software.broadinstitute.org/gsea/msigdb`).

| | **Gene set** | **Overlap** | **Genes in set** | $q$**-value** |
|---|---|---|---|---|
| 1 | protein localization | 215 | 1805 | 4.91E-55 |
| 2 | phosphate containing compound metabolic process | 226 | 1977 | 4.91E-55 |
| 3 | regulation of anatomical structure morphogenesis | 155 | 1021 | 2.1E-52 |
| 4 | positive regulation of molecular function | 208 | 1791 | 1.15E-51 |
| 5 | positive regulation of catalytic activity | 185 | 1518 | 1.14E-48 |
| 6 | regulation of multicellular organismal development | 190 | 1672 | 1.41E-45 |
| 7 | positive regulation of gene expression | 191 | 1733 | 6.24E-44 |
| 8 | positive regulation of biosynthetic process | 195 | 1805 | 1.1E-43 |
| 9 | small molecule metabolic process | 192 | 1767 | 2.19E-43 |
| 10 | regulation of transcription from rna polymerase ii promoter | 193 | 1784 | 2.19E-43 |
| 11 | cellular macromolecule localization | 156 | 1234 | 7.56E-43 |
| 12 | intracellular signal transduction | 177 | 1572 | 5.85E-42 |
| 13 | positive regulation of response to stimulus | 198 | 1929 | 2.68E-41 |
| 14 | lipid metabolic process | 147 | 1158 | 1.53E-40 |
| 15 | regulation of cell differentiation | 169 | 1492 | 2.3E-40 |
| 16 | positive regulation of cellular component organization | 146 | 1152 | 3.19E-40 |
| 17 | regulation of intracellular signal transduction | 179 | 1656 | 3.58E-40 |
| 18 | tissue development | 170 | 1518 | 4.79E-40 |
| 19 | single organism biosynthetic process | 157 | 1340 | 3.12E-39 |
| 20 | cellular response to organic substance | 189 | 1848 | 3.12E-39 |

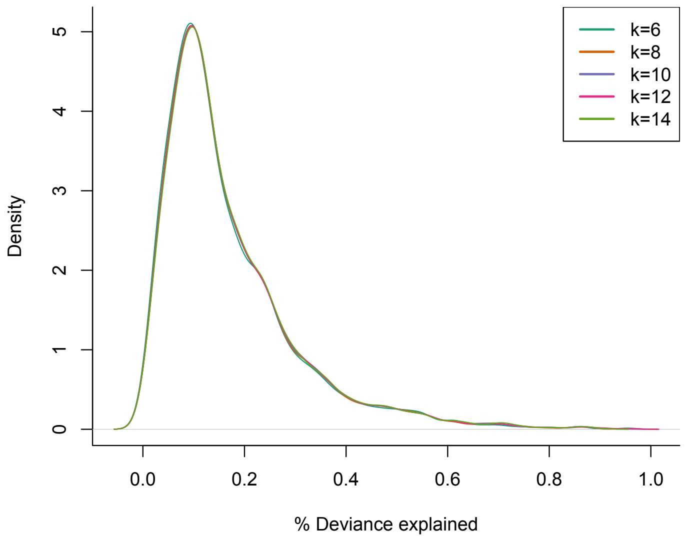


Figure S1: *Mouse bone marrow dataset: The NB-GAM is robust to the number of knots $k$.* Gaussian kernel density plot of the percentage of deviance explained by the NB-GAM model applied to each of the genes in the dataset from Paul et al. [2015], with number of knots $k$ ranging from 6 to 14. The distributions are nearly identical for the different numbers of knots.

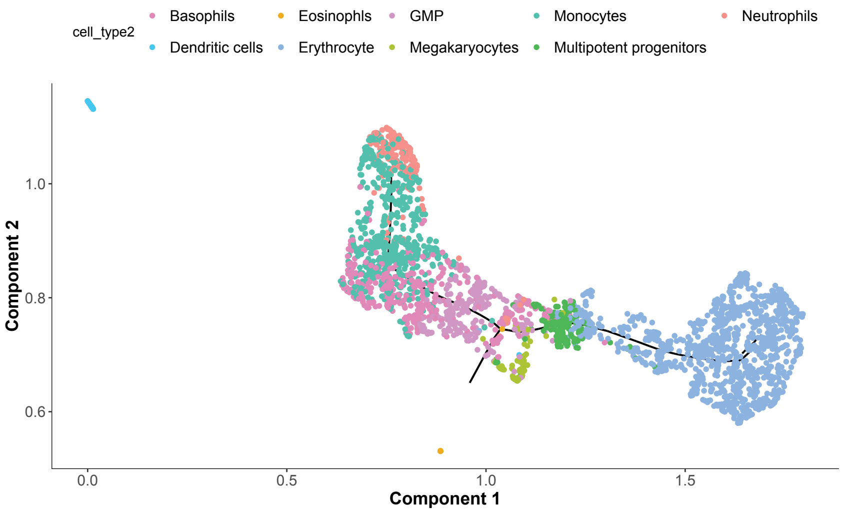


Figure S2: *Mouse bone marrow dataset: Outlying dendritic cells and eosinophils in UMAP space for TI with Monocle 3.*

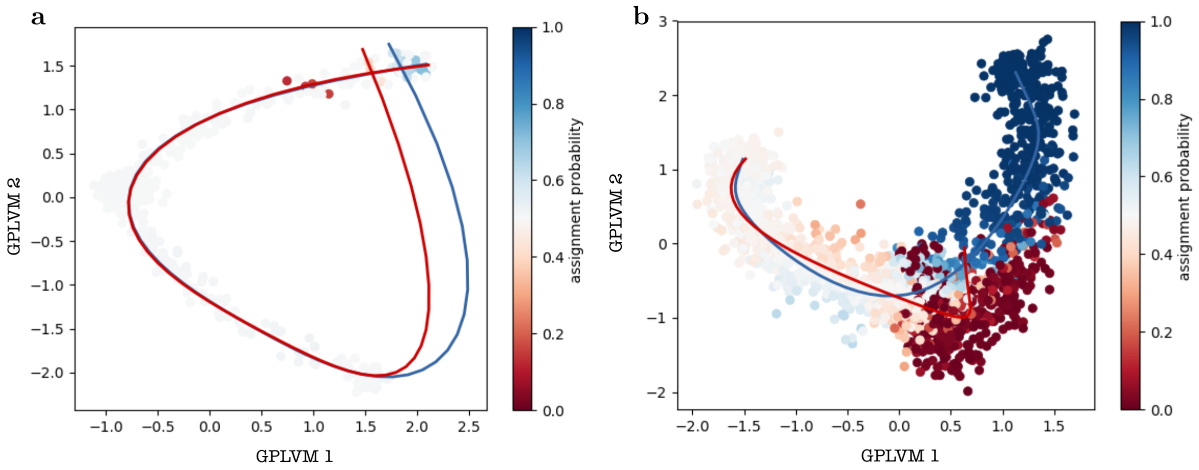


Figure S3: *Bifurcating simulation scenario: GPfates only recovers meaningful trajectories if the true pseudotime is provided as input.* Example of a bifurcating dataset from the dynverse framework. The dataset is represented in low-dimensional space using Gaussian latent variable models as implemented in GPfates. Cells are colored according to their assignment probability to a respectively colored lineage. Trajectories inferred by GPfates are shown when **(a)** pseudotime is estimated by GPfates and **(b)** true pseudotime is provided as input to GPfates.

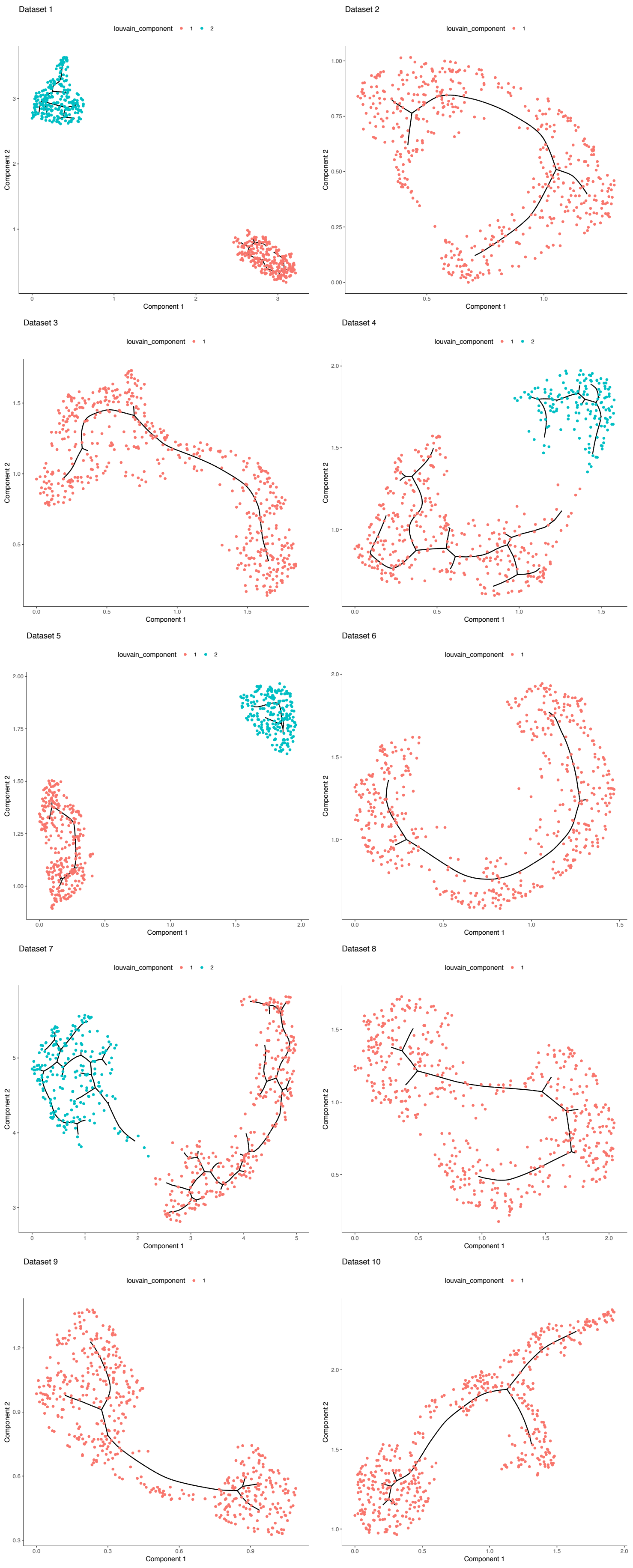


Figure S4: *Cyclic simulation scenario:* Monocle 3 *inferred trajectories for each of the* 10 *simulated datasets.* The first two components from UMAP dimensionality reduction, as implemented in Monocle 3, are plotted along with the Monocle 3 inferred trajectories. Cells are colored according to a Louvain clustering implemented in Monocle 3. Monocle 3 often fails to recover the cyclic pattern.

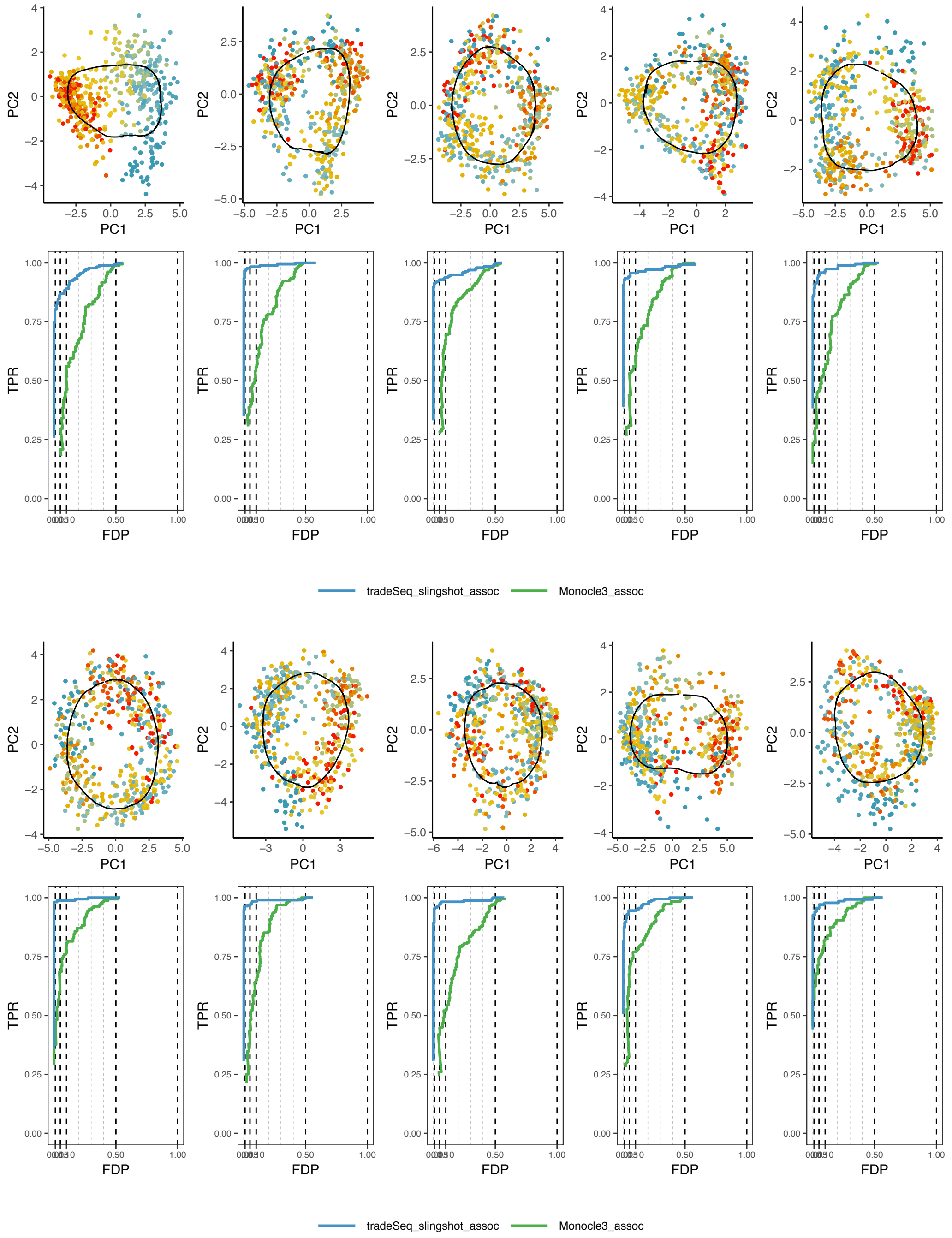


Figure S5: *Cyclic simulation scenario: FDP-TPR performance curves for trajectory-based differential expression analysis for each of the 10 simulated datasets.*

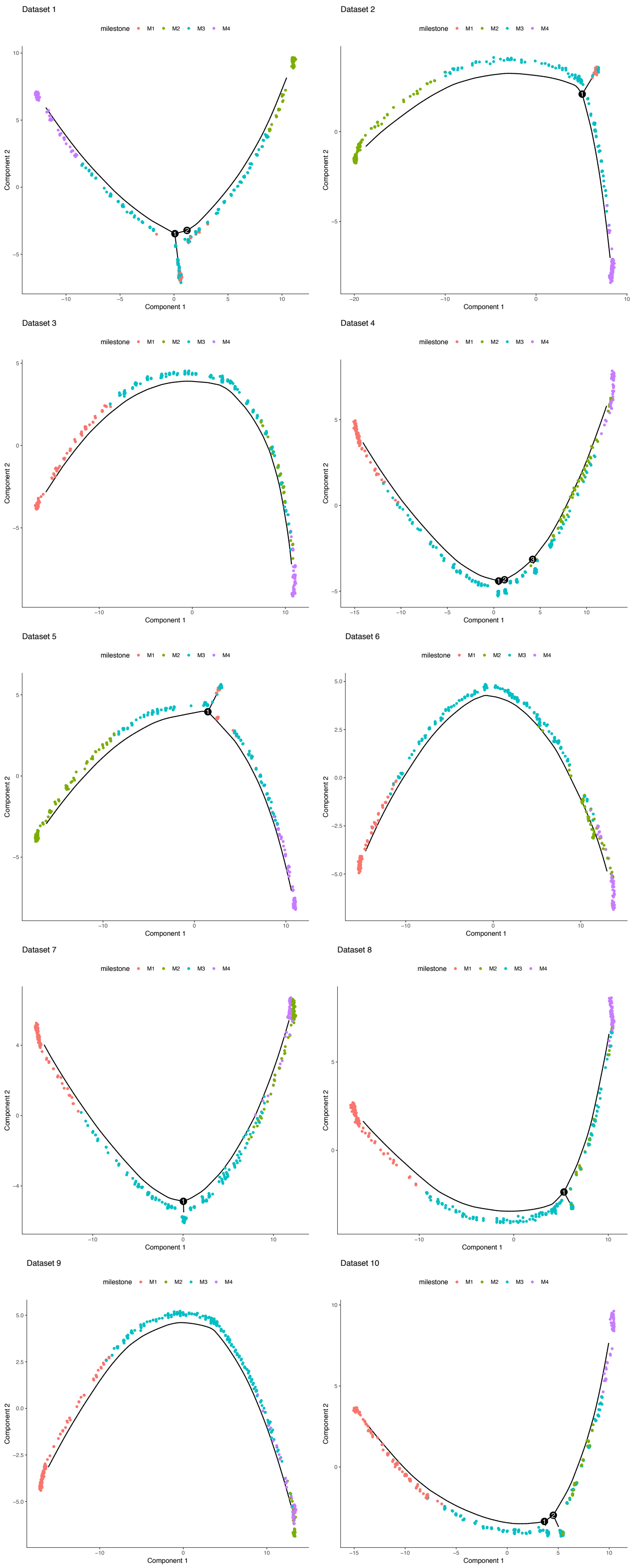


Figure S6: *Bifurcating simulation scenario: Monocle 2 inferred trajectories for each of the* 10 *the simulated datasets.* Cells are plotted in two-dimensional space using DDRTree dimensionality reduction [Qiu et al., 2017]. The simulated trajectory starts at milestone 1 and then continues into milestone 3, generating the two lineages that consist of milestone 2 and milestone 4. The trajectory is correctly recovered in, for example, Dataset 1 (top left panel). Dataset 4, on the other hand, wrongly assigns milestone 2 and milestone 4 to the same lineage, hence failing to recover the true bifurcation point.

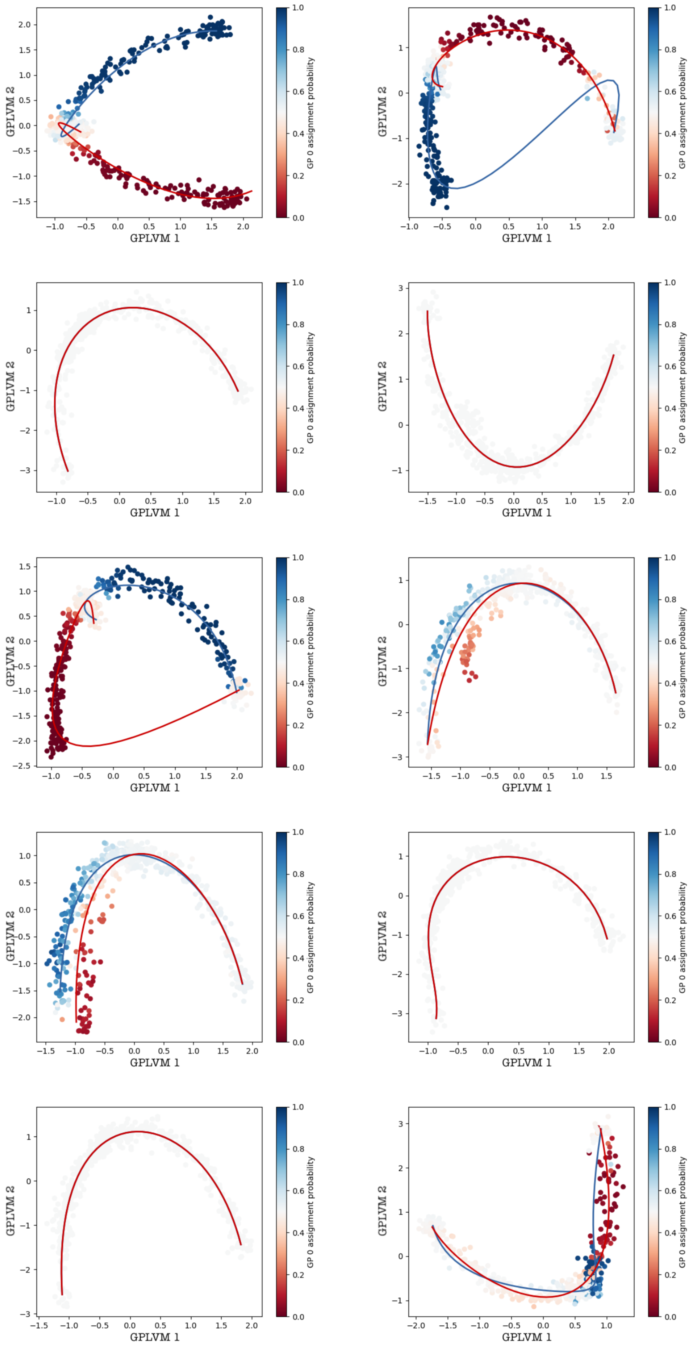


Figure S7: *Bifurcating simulation scenario:* GPfates *inferred trajectories for each of the* 10 *simulated datasets.* Two-dimensional representation of the datasets for the bifurcating simulation scenario (dynverse toolbox) using Gaussian latent variable models as implemented in GPfates. Cells are colored according to their assignment probability to a respectively colored lineage.

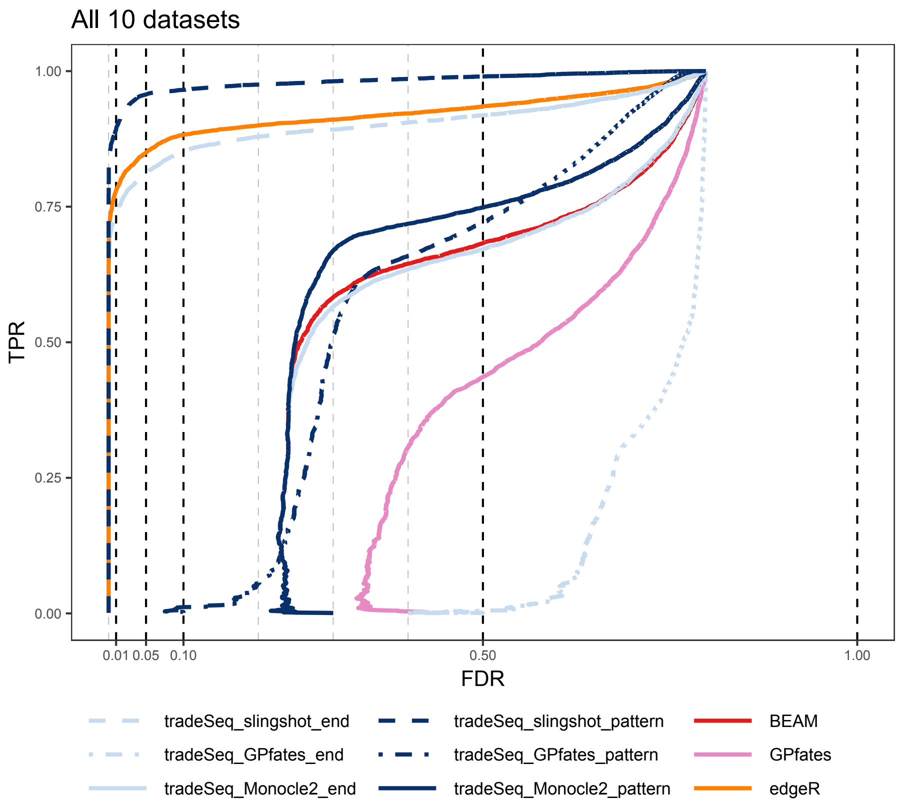


Figure S8: *Bifurcating simulation scenario: Mean FDR-TPR performance curves for trajectory-based differential expression analysis across all* 10 *simulated datasets.*

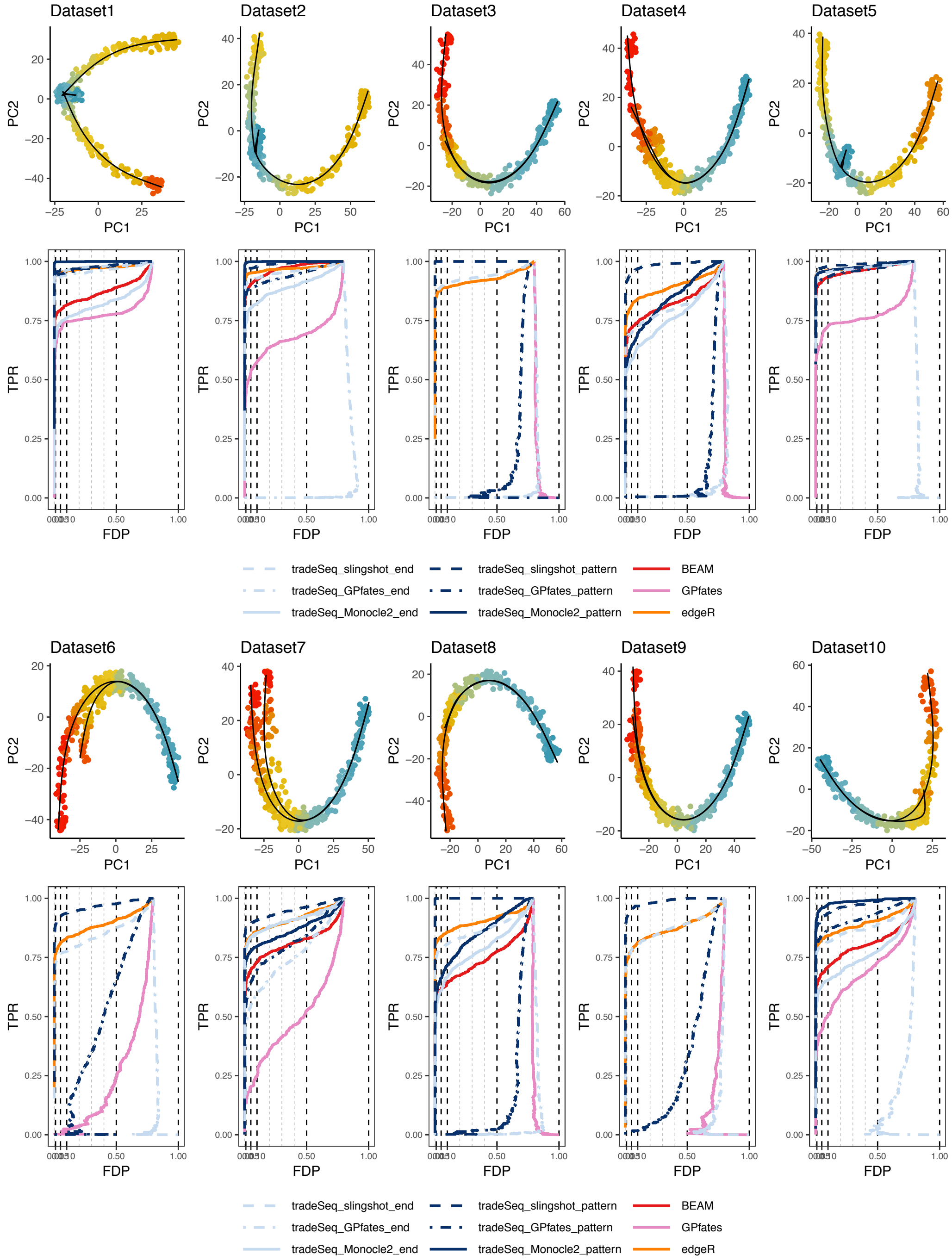


Figure S9: *Bifurcating simulation scenario: FDP-TPR performance curves for trajectory-based differential expression analysis for each of the* 10 *simulated datasets.* Note that the BEAM and tradeSeq_Monocle2 methods are not plotted for Datasets 3, 6 and 9, since Monocle2 failed to discover a branching trajectory for those datasets.

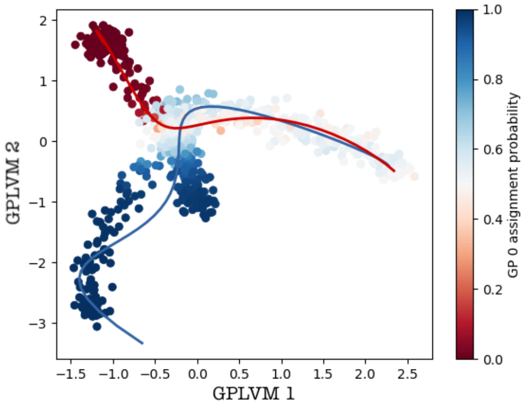


Figure S10: *Multifurcating simulation scenario:* `GPfates` *inferred trajectory on one simulated dataset.* Two-dimensional representation of the dataset for the multifurcating simulation scenario using Gaussian latent variable models, as implemented in `GPfates`. Cells are colored according to their assignment probability to a respectively colored lineage.

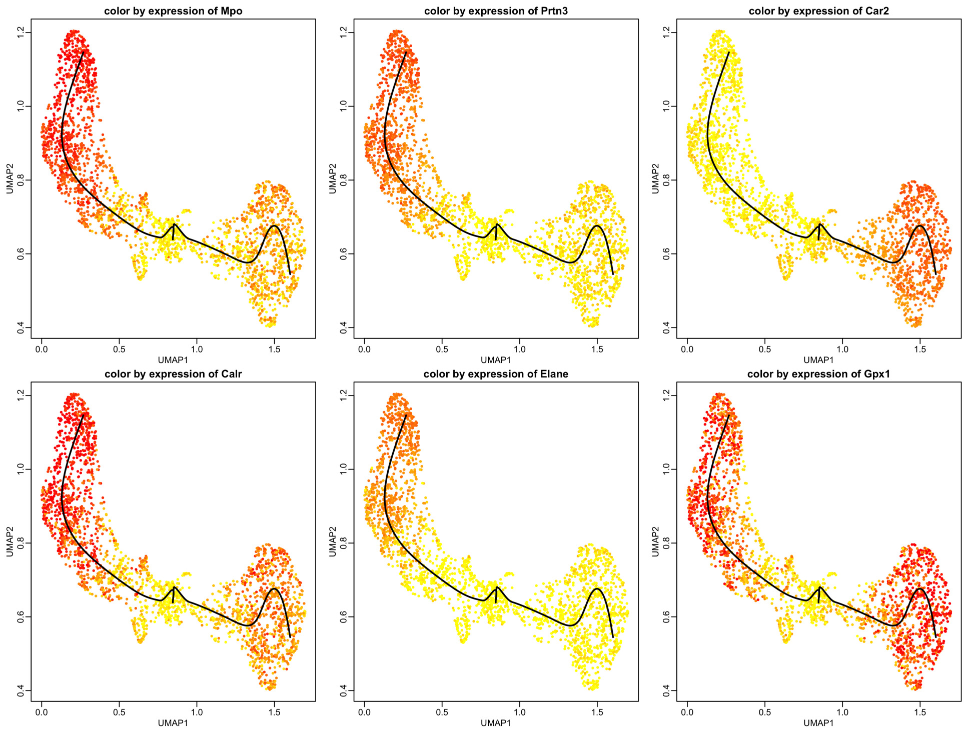


Figure S11: *Mouse bone marrow dataset:* `tradeSeq` *recovers markers for the progenitor cell population.* This figure shows the six most significant genes when testing for differential expression between the progenitor cell type (i.e., starting point of the smoother) and differentiated cell types (i.e., end point of the smoother) for the data from Paul et al. [2015] using `tradeSeq`. Yellow denotes low expression, while red denotes high expression.

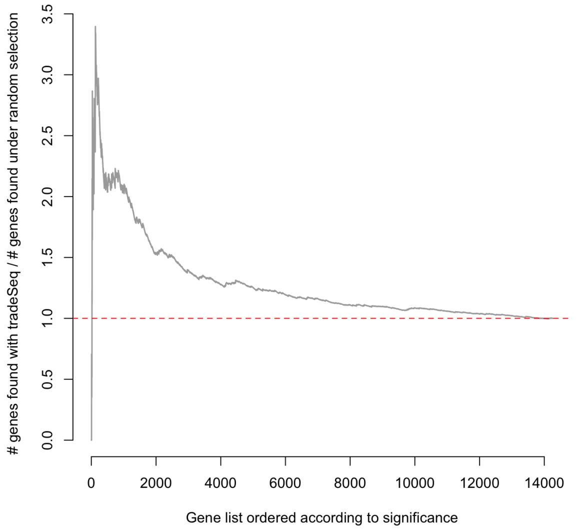


Figure S12: *Mouse olfactory epithelium dataset: Cell cycle genes in the neuronal lineage.* The figure illustrates the enrichment of cell cyle genes in lists of genes whose expression was found to be most significantly associated with the neuronal lineage according to the `associationTest` procedure in `tradeSeq`. The list of cell cycle related genes was obtained from the Mouse Genome Informatics (MGI) website at http://www.informatics.jax.org/go/term/GO:0007049. On the x-axis, genes are ordered according to their significance based on `associationTest`. The y-axis shows the ratio of the number of cell cycle genes among a set of top significant genes relative to the number of cell cycle genes one would expect by chance (i.e., if cell cycle genes were randomly found DE/sampled). If we let $C$ denote the proportion of genes associated with the cell cycle according to the MGI database, then, under the hypothesis that cell cycle genes are randomly discovered as DE, the expected number of cell cycle genes in the list of top $N$ genes is $NC$. The relative number that is plotted on the y-axis is then the ratio between the number of cell cycle genes discovered by `tradeSeq` for a given top list of size $N$ and $NC$.

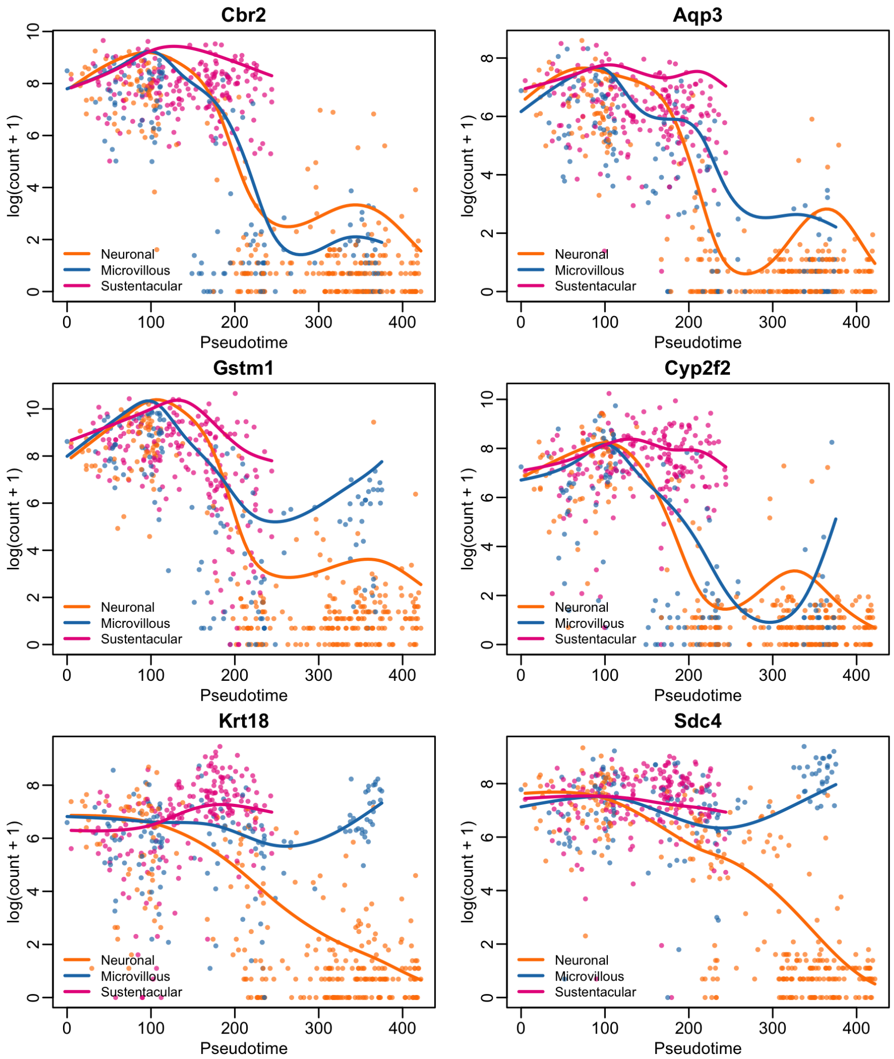


Figure S13: *Mouse olfactory epithelium dataset: Top six differentially expressed genes as identified by a ZINB analysis with* `tradeSeq` `patternTest`. Every trajectory is represented by a smooth function of gene expression along pseudotime. The cells assigned to a particular lineage based on the `slingshot` weights are represented with the same color as the lineage.

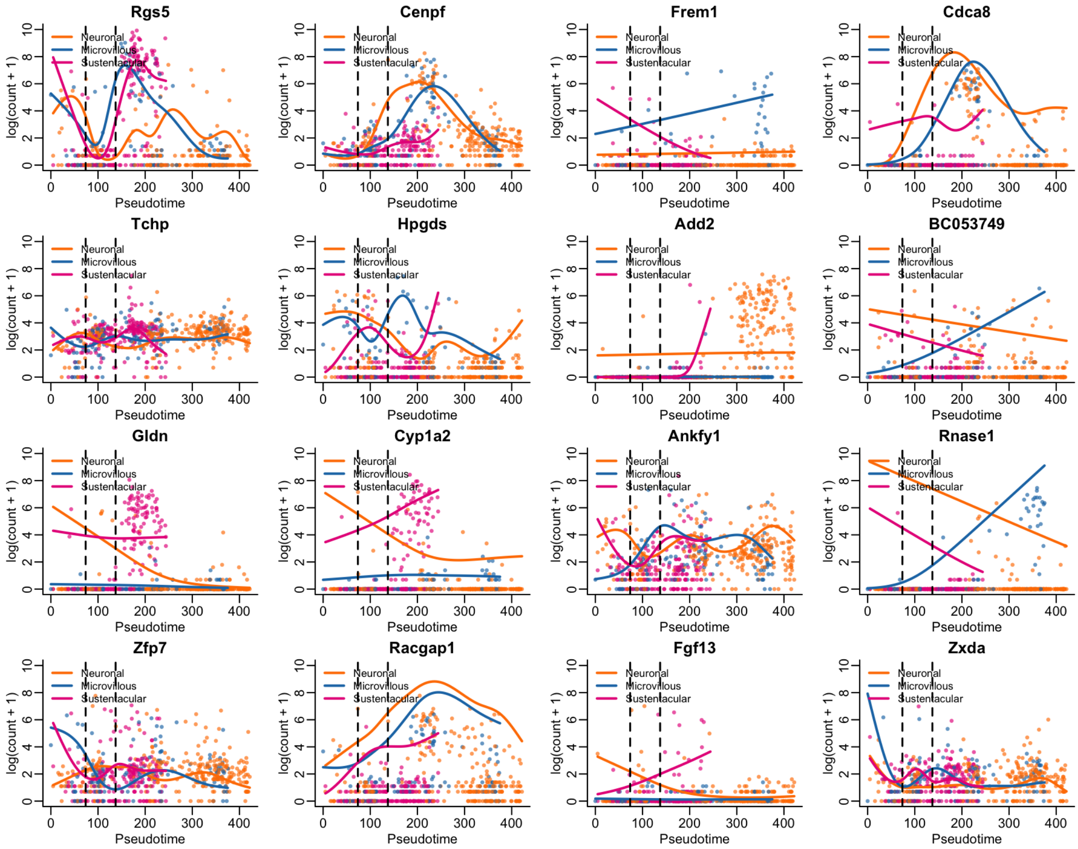


Figure S14: *Mouse olfactory epithelium dataset: All* 16 *genes found to be significant in all pairwise between-lineage comparisons based on a ZINB analysis with* `tradeSeq`*'s* `earlyDETest` *procedure over knots* 2 *and* 4. The cells assigned to a particular lineage based on the `slingshot` weights are represented with the same color as the lineage.

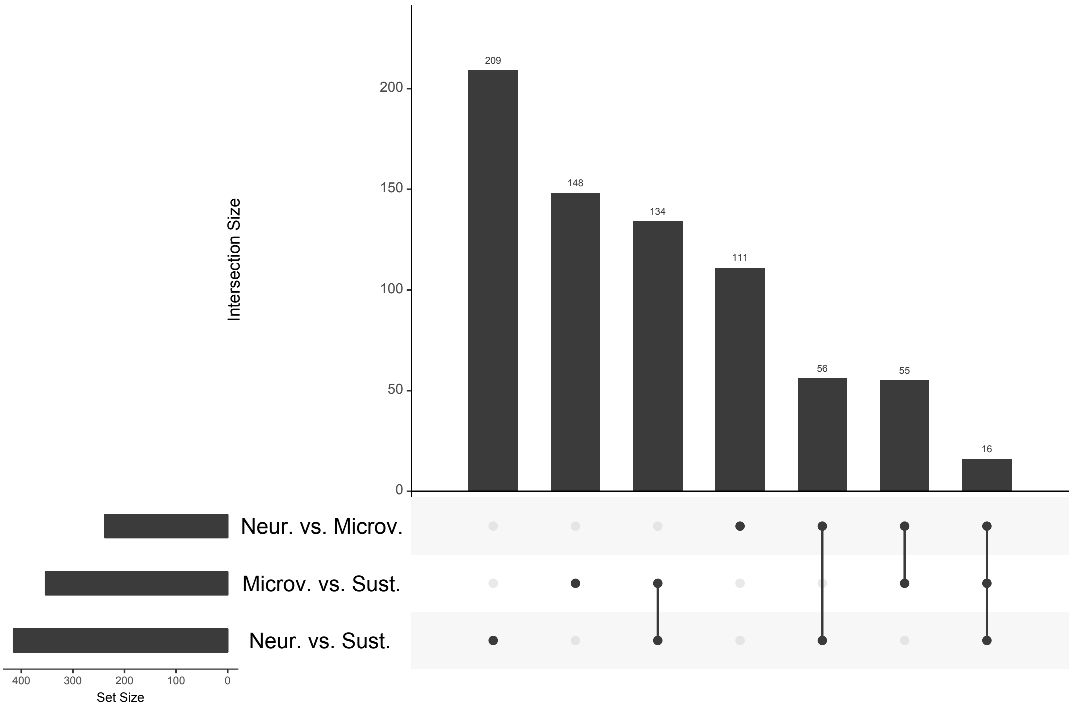


Figure S15: *Mouse olfactory epithelium dataset: UpSet plot for* `earlyDETest` *applied around the first branching point to each pair among the three lineages.* Left panel: Barplot of the number of DE genes for each contrast/pair (5% nominal FDR level). Top panel: Barplot of the number of DE genes for one, two of the three, or all three contrasts. Center panel: Contrasts being considered for the top panel. For example, the third column shows that 134 genes are DE for both the Microv vs. Sust contrast and the Neur. vs. Sust contrast.

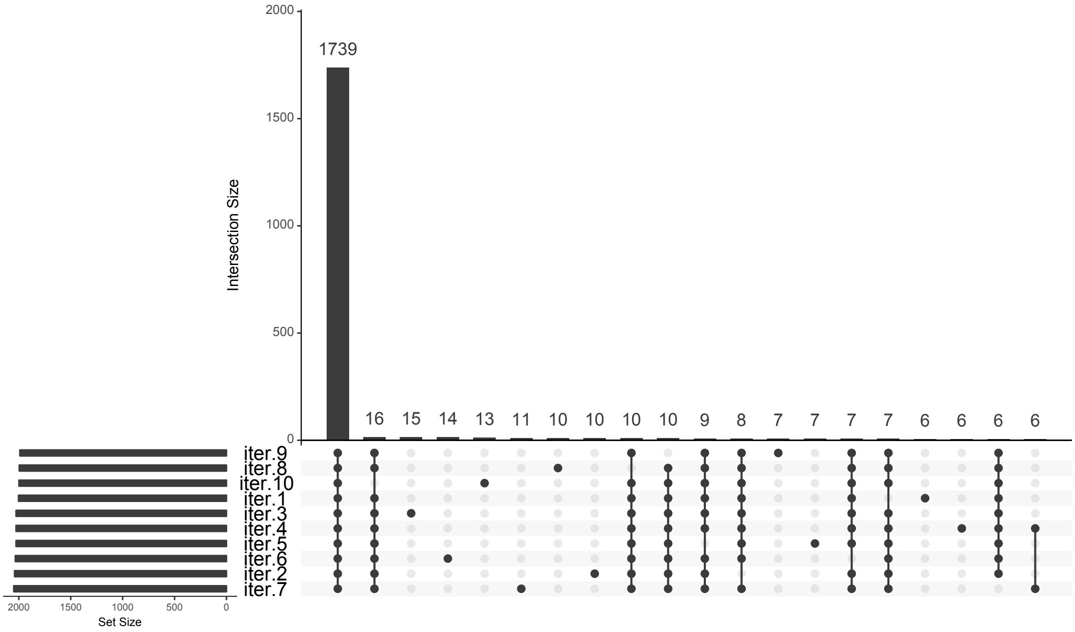


Figure S16: *Mouse bone marrow dataset: UpSet plot for* `startVsEndTest`*, for multiple assignments of cells to lineages.* Ten random multinomial assignments of cells to lineages are performed and denoted with ‘iter.1’ to ‘iter.10’. Left panel: Barplot of the number of DE genes for each assignment (5% nominal FDR level). Top panel: Barplot of the number of DE genes in common for various combinations of the 10 assignments. Center panel: Assignments being considered for the top panel. Only the 20 combinations with the largest number of common genes are shown in the figure.
